# Integration of individual and population data to improve predictions of structured population dynamics

**DOI:** 10.64898/2026.01.31.703022

**Authors:** Malcolm Itter

## Abstract

Integral projection models (IPMs) are a powerful tool for predicting structured population dynamics under global change. Inverse calibration approaches allow IPMs to be fit using widely available population density data increasing potential applications. Yet population density data alone may not provide sufficient information for IPMs to accurately identify the environmental factors driving demographic rates leading to poor predictions under variable future conditions. We construct a Bayesian dynamical IPM framework to integrate population density with individual growth data to better predict size-structured population dynamics. The framework pairs an IPM process model with data models that control for individual growth variability and a mismatch in the resolution and precision of population density and individual growth data. The model is applied to a combination of experimental and simulated forest population data to assess its ability to predict size-structured population density and estimate underlying demographic rates. Predictions of size-structured population density were similar regardless of whether the dynamical IPM was provided both population density and individual growth data (integrated model) or population density data alone (population model). The population model, however, did not identify annual growth effects driven by weather variables including vapor pressure deficit leading to poor estimates of mean size-structured growth. Simulation trials under which the integrated model was provided varying numbers of individual growth records indicated that 10 records were sufficient for the model to estimate annual growth effects with near equivalent inference when 30 or more records were applied. Results highlight the potential for inversely calibrated IPMs to correctly predict structured population dynamics while incorrectly estimating underlying demographic rates. Integrating individual demographic data resolves this issue allowing for inference on growth responses to variable environmental conditions, thereby improving the ability of IPMs to predict structured population dynamics under global change. While individual demographic rate data is often limited, simulation results indicate that only a small number of individual records are needed for valid inference.

## 1 Introduction

Global change is already contributing to fundamental shifts in ecosystem dynamics with impacts on biodiversity and ecosystem function (Moore and Schindler, 2022). Predictions of ecosystem dynamics under global change are needed to help inform conservation management (Yates et al., 2018). Population structure (e.g., distribution of ages, sizes, functional traits) is important in this context because it may support ecosystem resistance, resilience, and adaptive capacity to novel conditions (Nicotra et al., 2015). Integral projection models (IPMs) are a well-known class of models to predict structured population dynamics while allowing for inference on demographic responses to environmental conditions (Ellner and Rees, 2006). Yet constructing well-calibrated IPMs capable of making transferable predictions along with estimates of associated uncertainty is challenging given the need for large-scale spatio-temporal data that contains sufficient information about population-level responses to novel conditions (Gelfand et al., 2013).

The growth, mortality, and regeneration functions that underlie an IPM are traditionally fit to individual demographic data using generalized linear models (Rees et al., 2014). Parameterized demographic functions are then plugged into a broader IPM framework to predict population dynamics. An alternative is to apply an inverse calibration approach under which observations of structured population density over time are used to infer underlying demographic rates and their responses to the environment (Ghosh et al., 2012; González et al., 2016). Inverse calibration requires a probabilistic model framework to connect latent (unobserved) demographic rates to noisy, population-level observations of their outcomes. Bayesian state space frameworks are well suited to this problem and have been used in several studies applying an inverse calibration approach with the structured population density representing the latent state and the IPM serving as the process model (Ghosh et al., 2012; White et al., 2016; Plard et al., 2019b; Shriver et al., 2021).

Integrating an IPM as a process model within a broader Bayesian state space framework (hereafter referred to as a “Bayesian dynamical IPM”) allows the IPM to be fit when only population data is available and can accommodate missing observations in the form of irregular population density time series thereby broadening their applicability (White et al., 2016). Bayesian dynamical IPMs support deeper understanding of potential demographic constraints on population dynamics through estimated correlation structures among jointly modeled demographic parameters (González et al., 2016). Further, Bayesian dynamical IPMs facilitate propagation of uncertainty in underlying demographic functions, model parameters, and noisy observations to predictions of emergent population dynamics (Elderd and Miller, 2016). Lastly, such models allow for the integration of multiple datasets with potentially different scales (e.g., sampling unit, temporal resolution of observations, level of precision) to inform parameter estimates and predictions of structured population dynamics (Plard et al., 2019a).

Despite the benefits of a Bayesian dynamical IPM, it can be challenging to infer demographic rates from population data alone. Under this approach, the model must separate and identify the factors affecting multiple demographic rates using only noisy observations of their emergent outcomes (González et al., 2016). These models may suffer from equifinality under which multiple demographic parameter combinations lead to similar predictions of population dynamics. For example, an IPM may correctly predict structured population density over time while incorrectly estimating the factors driving underlying demographic rates. This may lead to poor predictions of future population dynamics for which there are no observations to constrain the process, particularly if the environmental variables driving demographic rates undergo fundamental changes in the future (Fig. 1).

**Figure 1:**
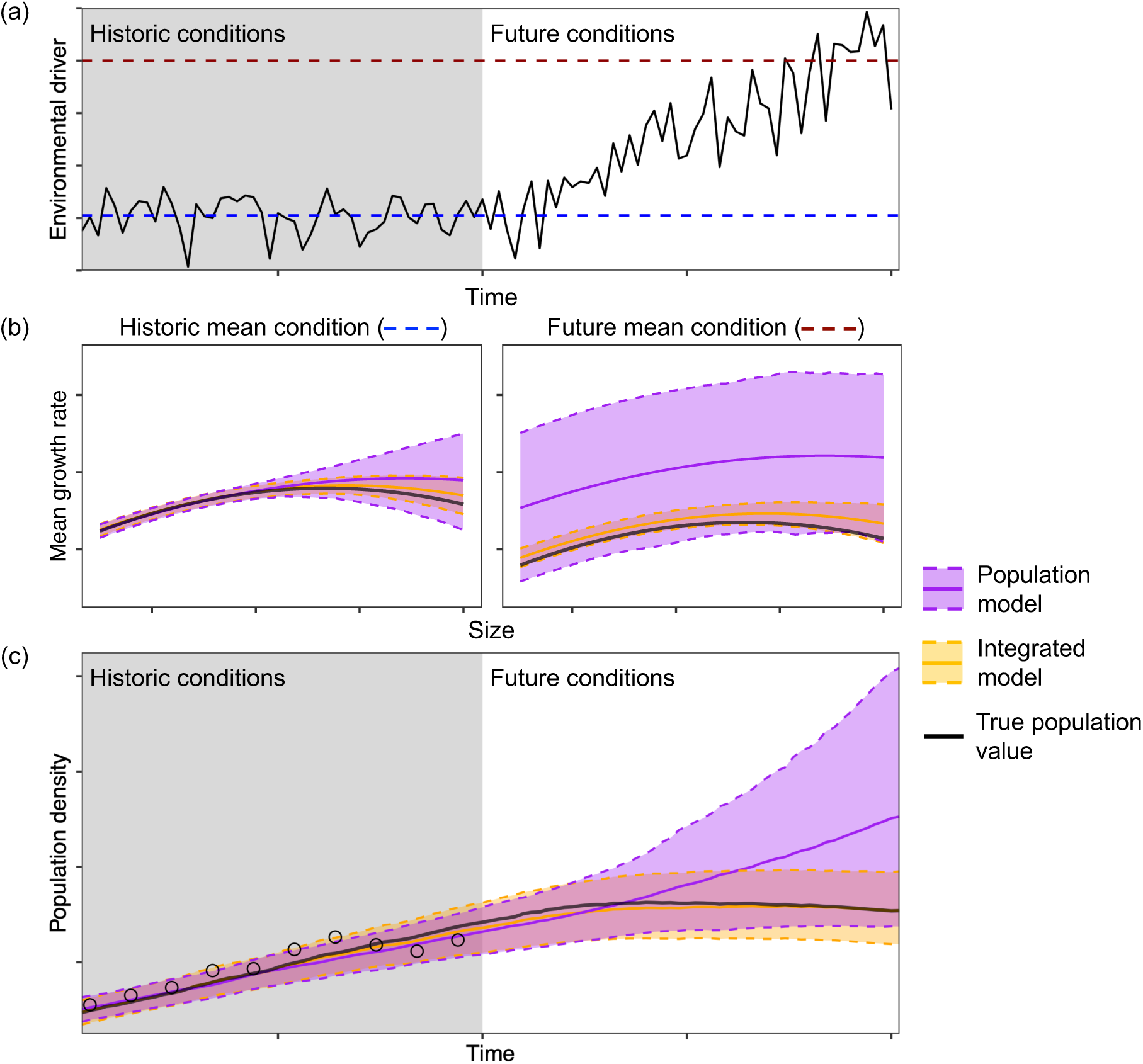
Example predictions of size-structured population density under historic and future environmental conditions applying population density data alone (population model) and a combination of population density and individual growth rate data (integrated model). Panel (a) depicts change in the environmental driver over time highlighting increasing mean and variance under future conditions. Panel (b) depicts modeled mean size-structured growth under the mean historic condition (blue line) and the mean future condition (red line). Panel (c) depicts predictions of size-structured density applying the mean growth estimates in (b) with points representing periodic population density observations.

Previous Bayesian dynamical IPM applications have used informed prior distributions to ensure the idenfiability and separability of important demographic parameters (White et al., 2016), but this requires previous understanding of the factors driving demographic processes. Another approach is to integrate population and individual demographic data leveraging the flexibility of Bayesian dynamical IPMs to accommodate multiple data sources, building on well-developed integrated population model methodology (Zipkin and Saunders, 2018; Plard et al., 2019a). The goal of such data integration is to better identify the factors driving demographic rates while accurately predicting structured population dynamics under future environmental conditions (Fig. 1). For example, Plard et al. (2019b) constructed an integrated Bayesian dynamical IPM that correctly identified the relationships between environmental variables and individual demographic rates, while accurately predicting structured population dynamics over time.

Integrating structured population density and individual demographic data within a Bayesian dynamical IPM presents several modeling challenges. Specifically, individual demographic rates are strongly affected by individual variability associated with often unobserved variables such as genotype, micro-habitat, local competition, and functional traits (Clark et al., 2010). This variability may bias estimates of the factors driving mean demographic rates within the modeled population (Ghosh et al., 2012). Further, individual demographic rate observations may exhibit strong temporal autocorrelation particularly if they are related to time-varying environmental conditions such as shown in Figure 1 (Fay et al., 2022). Lastly, individual demographic data may be misaligned with population data both in terms of its temporal resolution and precision.

We present an integrated Bayesian dynamical IPM aimed at addressing the above modeling challenges in a generalizable manner. The model connects population density and individual growth rate data observed at different temporal scales and precisions to an underlying IPM process model describing changes in structured population density over time. It accounts for individual variation in observed growth rates attributable to both observed and unobserved individual variables. We apply the Bayesian dynamical IPM to model size-structured forest population dynamics using population data only (population model) and a combination of population and individual growth rate data (integrated model). We compare predictions of structured population dynamics and estimates of underlying mean growth rates under variable environmental conditions over time using a combination of simulated and experimental forest data. Simulated data is used to assess how inference changes as a function of the number of individual growth rate observations used to inform the dynamical IPM.

We focus model application on forest population dynamics because they highlight many of the previously described data challenges related to integration of population density and individual demographic rate data (Evans et al., 2022). Further, inverse calibration approaches are increasingly used to calibrate models of forest population dynamics including IPMs (Ghosh et al., 2012) and broader dynamic vegetation models (Hartig et al., 2012). While the model application is specific to forest population dynamics, the methods are applicable to a broad diversity of ecological data.

## 2 Materials and methods

### 2.1 Data

In the context of modeling forest population dynamics, observations of size-structured population density are provided by forest inventory data consisting of repeat measures of individual tree size, species, and status (live/dead) within permanent sample plots on a periodic basis (e.g., every 5-10 years). Observations of individual growth come from tree ring data providing measures of diameter growth at breast height (1.3 m) on an annual basis for a subset of individuals from the population. Periodic inventory data makes it challenging to infer demographic responses to environmental variables such as weather or disturbance that vary greatly on an inter-annual basis. Annual observations of individual growth provide rich information about environmental drivers, but are strongly affected by previously described individual growth variability.

#### 2.1.1 Simulated data

Simulated data was used to assess the ability of the Bayesian dynamical IPM to predict structured population density over time and infer associated demographic parameters based on different amounts of population density and individual growth data. Data were simulated for a single species from the proposed dynamical IPM (Section 2.2) for a 50-year period. Population density data consisted of inventory observations for ten 0.1 ha forest inventory plots generated annually or periodically at five-year increments providing observations of diameter at breast height (DBH) for all live trees with DBH ⩾ 10 cm within each plot. Individual data consisted of annual diameter growth rate observations for up to 200 individuals co-located with the simulated inventory plots. A single time varying environmental variable representing annual mean monthly vapor pressure deficit (VPD) during summer months (June - August) was simulated as a driver of growth.

#### 2.1.2 Birch Lake red pine site

Birch Lake is a USDA Forest Service experimental forest site in northeastern Minnesota, USA (47*^◦^*43’N, 91*^◦^*55’W). The experimental site was established in 1957 in a 45-year-old red pine (*Pinus resinosa* Ait.) plantation to assess forest demographic and structural responses to alternative thinning methods and intensities (Powers et al., 2010). The Birch Lake site offers a unique combination of long-term inventory and tree ring data from a large sample of co-located red pine individuals. Forest inventory plots 0.08 ha in size were established within each Birch Lake experimental unit in 1957 (one plot per unit). All trees with DBH ⩾ 10 cm were tagged within each plot and their DBH measured to the nearest 0.3 cm (0.1 in) using diameter tape. We applied data collected within nine forest inventory plots located within control units receiving no management intervention. Each plot was measured ten times between 1957-2009 (additional detail on the Birch Lake study design and inventory data collection is provided in Powers et al. (2010)). Increment cores were collected at breast height from all live trees with DBH ⩾ 5 cm within each inventory plot in 2009. All increment cores were processed and cross-dated using standard methodologies providing observations of radial growth increment on an annual basis (D’Amato et al., 2013). There were a total of 652 trees cored across the nine control plots. Select monthly weather variables including maximum monthly VPD and temperature (T^Max^) were extracted for the Birch Lake site for years 1957-2009 from the Parameter-elevation Regressions on Independent Slopes Model (PRISM Group, Oregon State University, 2025). Monthly weather variables were converted into annual mean monthly estimates of summer VPD (June-August) and spring T^Max^ (March-May), which were applied to model annual growth rates.

### 2.2 Dynamical IPM framework

We apply a Bayesian hierarchical approach to predict size-structured population dynamics over time integrating population density and individual growth rate data (Berliner, 1996). The hierarchical framework includes data models for population density and individual growth observations, an IPM process model, and models for unknown data and process parameters (Figs. 2, S1; Table S1).

**Figure 2:**
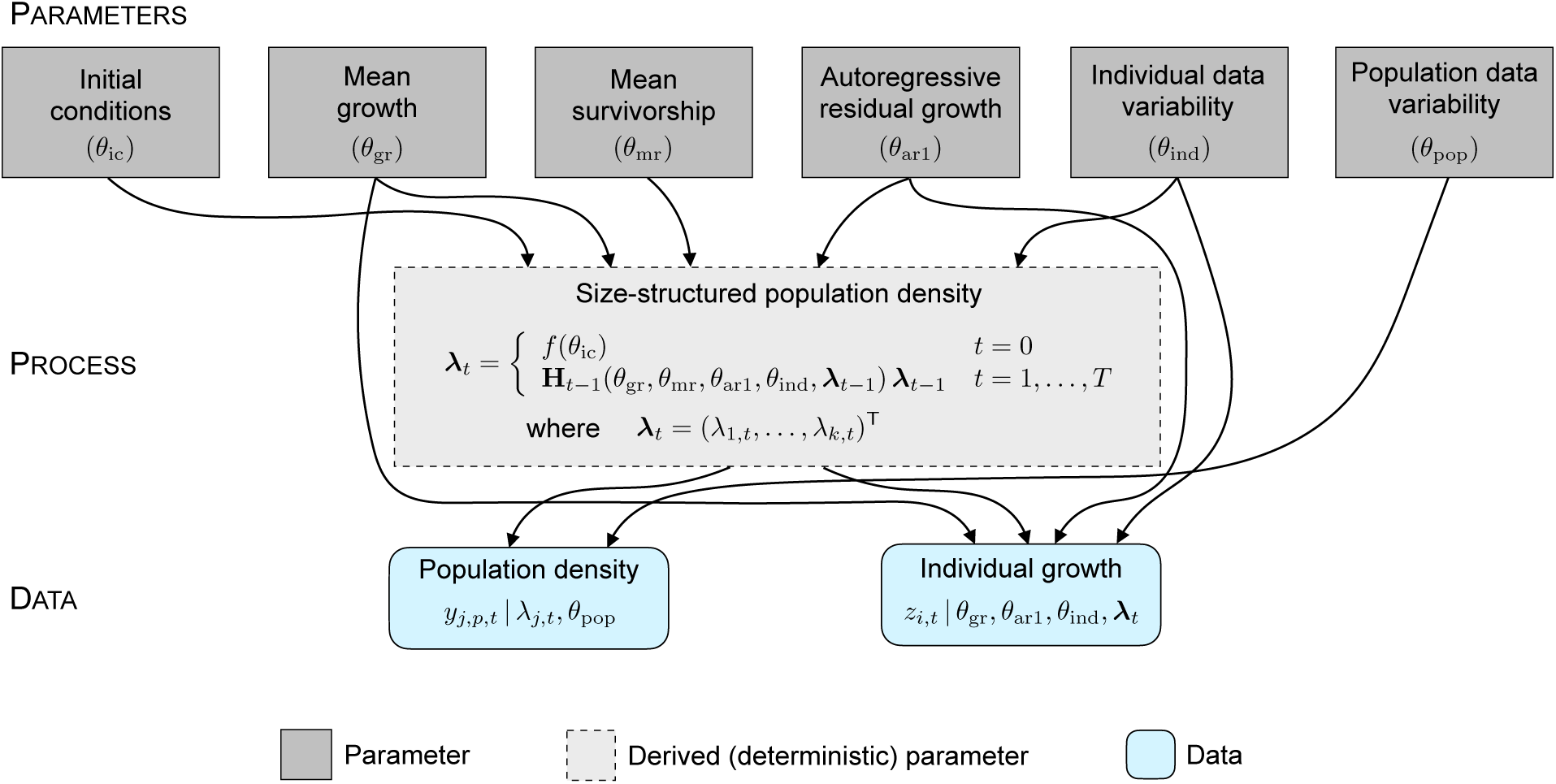
Graphical representation of the integrated dynamical integral projection model (IPM) structure and parameter dependencies. Parameters are grouped by model component and defined in Table S1. The modeled process of interest is the latent size-structured population density. Throughout, *|* is used to denote conditional dependence.

#### 2.2.1 Data models

##### Population density data

Population density data include observations of the number and DBH (size) of individual trees within forest inventory plots. These observations represent a realization from an inhomogeneous Poisson point process in size generated conditional on the latent size-structured population density (Ghosh et al., 2012). Let *y_j,t,p_* denote the count of trees in size class *j* (*j* = 1*, …, k*) in plot *p* (*p* = 1*, …, P*) in inventory year *t* (*t* = 0*, …, T*), and *λ_j,t_* denote the latent mean population density in the corresponding size class. Population density observations are modeled applying a negative binomial likelihood

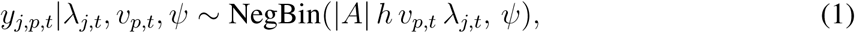

where *A* is the known size of the permanent sample plot in ha, *h* is the known size class width in cm, *v_p,t_* is a temporal plot random effect, and *ψ* is an overdispersion parameter. The plot random effect (*v_p,t_*) accounts for plots with consistently higher or lower observed densities than the mean population condition while the overdispersion parameter (*ψ*) allows for increased variability in observed counts.

#### Individual growth data

Individual growth rate data consist of tree ring records providing observations of annual DBH (size) growth increment for a selection of individuals from the population. Let *z_i,t_* denote the DBH growth in cm of the *i*th tree (*i* = 1*, …, n*) in year *t*. The observed growth period for an individual is unique and does not necessarily span the entire period over which population density is observed (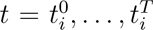 where 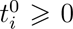 and 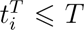). The data model connects observations of individual growth to the mean size-structured growth rate for the cohort of individuals of a given size *x* denoted as *δ_t_*(*x*). The aim of the individual growth data model is to allow the IPM process model to learn about the factors affecting mean size-structured growth (*δ_t_*(*x*)) while controlling for individual variation and temporal autocorrelation that may confound these effects. To this end, an individual random effect (*w_i_*) is used to account for consistent growth differences among individuals due to unobserved variables (e.g., genotype, micro-habitat, etc.) estimated conditional on an individual growth variance parameter 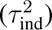. A first order autoregressive error structure is applied to account for temporal autocorrelation within individual growth records defined by an autoregressive coefficient (*ρ*) and white noise variance *σ*^2^. Individual growth observations are modeled applying a truncated normal likelihood

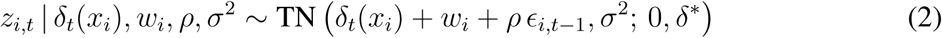

where *ɛ_i,t−_*_1_ is the residual error (*ɛ_i,t_* = *z_i,t_ −* (*δ_t_*(*x_i_*) + *w_i_*) for 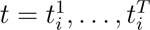. Growth observations are bounded between 0 and a fixed maximum potential growth rate *δ^∗^*.

#### 2.2.2 IPM process model

The process of interest is the latent size structured population density *λ_t_*(*s*). In the applied forest population data, *λ_t_*(*s*) expresses the number of trees per ha (density) in year *t* (*t* = 0, 1*, …, T*) of a given size *s* measured in terms of DBH. An IPM process model is applied to predict annual changes to the size structured population density,

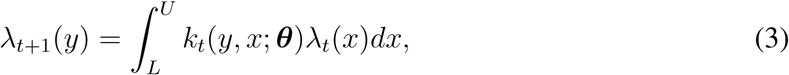

where *k_t_*(*y, x*; ***θ***) is a redistribution kernel describing the transition from size *x* to *y* in year *t* conditional on demographic parameters ***θ***, and *L*, *U* are the lower and upper size bounds of the modeled population. The IPM kernel consists of a growth kernel (*g_t_*(*x, y*)) modeling transition from size *x* to *y* and a mean size-structured survivorship term for the cohort of individuals of size *x* (*ξ_t_*(*x*)). While an IPM kernel generally also includes a regeneration term (Rees et al., 2014), we exclude it from the current model because our focus is on changes in size-structured density of a single demographic cohort (closed population that regenerated at approximately the same time with no new regeneration) as represented by the simulated and Birch Lake datasets.

A truncated normal growth kernel is applied to model the transition from size *x* to *y* conditional on the mean size-structured growth rate (*δ_t_*(*x*)) and variance *ν*^2^. As in the individual growth data model, growth is bounded to be between 0 and *δ^∗^*. Mean size-structured growth (*δ_t_*(*x*)) is estimated as a linear function of size, population density, weather conditions, and an annual random effect that accounts for additional inter-annual growth variation (Table 1). Mean size-structured survivorship (*ξ_t_*(*x*)) is estimated as a logistic function of size and population density (measured in terms of basal area per ha). The variance of the growth kernel (*ν*^2^) is calculated as the sum of the individual growth variance (*τ* ^2^) and the variance of the first order autoregressive residual error process (*σ*^2^(1−*ρ*^2^)−^1^) as defined in the individual growth data model (Fig. 2).

**Table 1:** Decomposition of mean size-structured growth into estimated fixed and random effects (DBH = diameter at breast height, BAH = basal area per ha, VPD = vapor pressure deficit, T^MAX^ = maximum temperature).

| Term | Expression | Description |
| --- | --- | --- |
| Intercept | $\beta_0$ | Mean growth rate integrating over time and size |
| Size | $\beta_1 \text{DBH} + \beta_2 \text{DBH}^2$ | Non-linear growth in relation to size |
| Population density | $\beta_3 \text{BAH}_t$ | Competitive effects of size-structured density |
| Weather* | $\beta_4 \text{VPD}_t + \beta_5 \text{VPD}_{t-1} + \beta_6 T_t^{\text{MAX}}$ | Environmental growth factors |
| Annual random effect | $u_t$ | Residual inter-annual growth variation |

\*Weather expression for simulated data includes $\beta_4 \text{VPD}_t$ only

The latent size-structured population density is modeled on a discrete scale applying a bin-to-bin method to numerically approximate the integral in Eqtn. (3) (Dawson, 2013; Ellner et al., 2016). The applied process model is expressed as

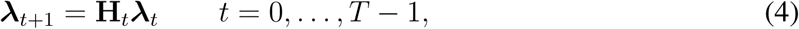

where ***λ****_t_* is a *k*-dimensional vector containing the mean population density for *k* equally-spaced discrete size classes (***λ****_t_* = (*λ*_1_*_,t_, …, λ_k,t_*)^T^), and **H***_t_* is a *k × k* propagator matrix that follows from the applied growth kernel, mean survivorship model, and the numerical integration method (Eqtn. S3).

#### 2.2.3 Parameter models and implementation

Unknown model parameters include size-structured mean growth and survivorship parameters, growth variance parameters, individual growth random effects, population density data parameters, and parameters representing the initial size distribution of the population (Fig. 2, Table S1). Models and prior distributions for all unknown parameters are included in the Supporting information. The dynamical IPM was fit using Markov chain Monte Carlo (MCMC) to sample from the joint posterior distribution. We applied a Hamiltonian Monte Carlo algorithm written in STAN (Stan Development Team, 2024) and implemented using the cmdstanr package (Gabry et al., 2025) for the R statistical computing environment (R Core Team, 2024). For all applications of the model, three MCMC chains were run for a total of 2,000 iterations with a burn-in of 1,000 iterations. Model convergence was assessed based on R-hat values (*<* 1.01), bulk effective sample size (minimum of 100 per chain), and visual inspection of chains for all parameters (Vehtari et al., 2021).

### 2.3 Model evaluation

The Bayesian dynamical IPM was fit using population density data alone (population model) and to the combination of population density and individual growth data (integrated model) to evaluate differences in the prediction of size-structured population density and inference on underlying mean growth rate parameters.

#### Simulated data

Seven simulated data scenarios were defined to evaluate different combinations of population density and individual growth data. Two population model scenarios were defined by fitting the dynamical IPM to either annual or periodic population density data alone, hereafter referred to as the annual population and periodic population model scenarios, respectively. Five integrated model scenarios paired periodic population density data with varying numbers of individual growth records (10, 30, 50, 100, 200). The annual population model scenario was used to assess whether potential differences in inference under integrated model scenarios are due to the annual resolution of individual growth data (as opposed to periodic population density data) or to other factors. For all simulation scenarios, the root mean square error (RMSE) was used to assess estimates of mean size-structured growth rate, and total population density in terms of stems and basal area per ha comparing simulated parameter values to model-based estimates. We further assessed recovery of simulated growth parameters based on 95 percent credible intervals.

#### Birch Lake data

We conducted 3-fold cross validation for the Birch Lake dataset to compare the ability of the population versus integrated models to predict out-of-sample size-structured population density and individual tree growth data. Within each fold, data were held out from three of the nine Birch Lake forest inventory plots (validation set) with the dynamical IPM fit using data from the remaining six plots (training set). Model predictions were evaluated using the log pointwise predictive density (lppd) score averaged across cross-validation folds (Gelman et al., 2014). The RMSE of total population density in terms of stems and basal area per ha and mean size-structured growth was also calculated. Variance partitioning was applied to determine the proportion of mean size-structured growth rate variation attributable to annual growth effects calculated as the sum of weather and annual random effects (Schulz et al., 2025), and the proportion of annual growth effect variability attributable to weather effects alone (Gelman et al., 2019). Additional information on model evaluation is provided in the Supporting information.

## 3 Results

### 3.1 Simulated data

Predictions of size-structured population density were roughly equivalent across simulated data scenarios (Table S2, Fig. S2). The RMSE of trees and basal area per ha derived from the latent population density were small for all scenarios (mean relative error ~ 1-2 percent) and generally decreased for integrated model scenarios with larger numbers of individual growth records (Table S2). In contrast, the ability of the dynamical IPM to accurately estimate growth rate parameters depended on the type and amount of simulated data used to inform it.

While the periodic population model accurately identified growth parameters associated with size and population density (Fig. S3), it did not well identify the single weather growth coefficient for VPD (*β*_4_) or the associated annual random effect standard deviation (*τ*_yr_) (Fig. 3). Specifically, the posterior distributions for these parameters were centered far away from simulated values with high levels of uncertainty resulting in large RMSE for estimates of mean size-structured growth (Fig. 3). Posterior estimates of annual growth effects showed limited inter-annual variation and frequently were in the wrong direction (posterior mean was positive when simulated effect was negative or vice versa; Fig. 4). There were also divergent transitions under the periodic population model (approximately 1-3 percent of post burn-in iterations resulted in a divergent transition). Providing the population model annual rather than periodic forest inventory data (annual population model) eliminated divergent transitions, and improved posterior inference for weather and annual random effects leading to more accurate and precise mean growth rate estimates (Figs. 3, 4).

**Figure 3:**
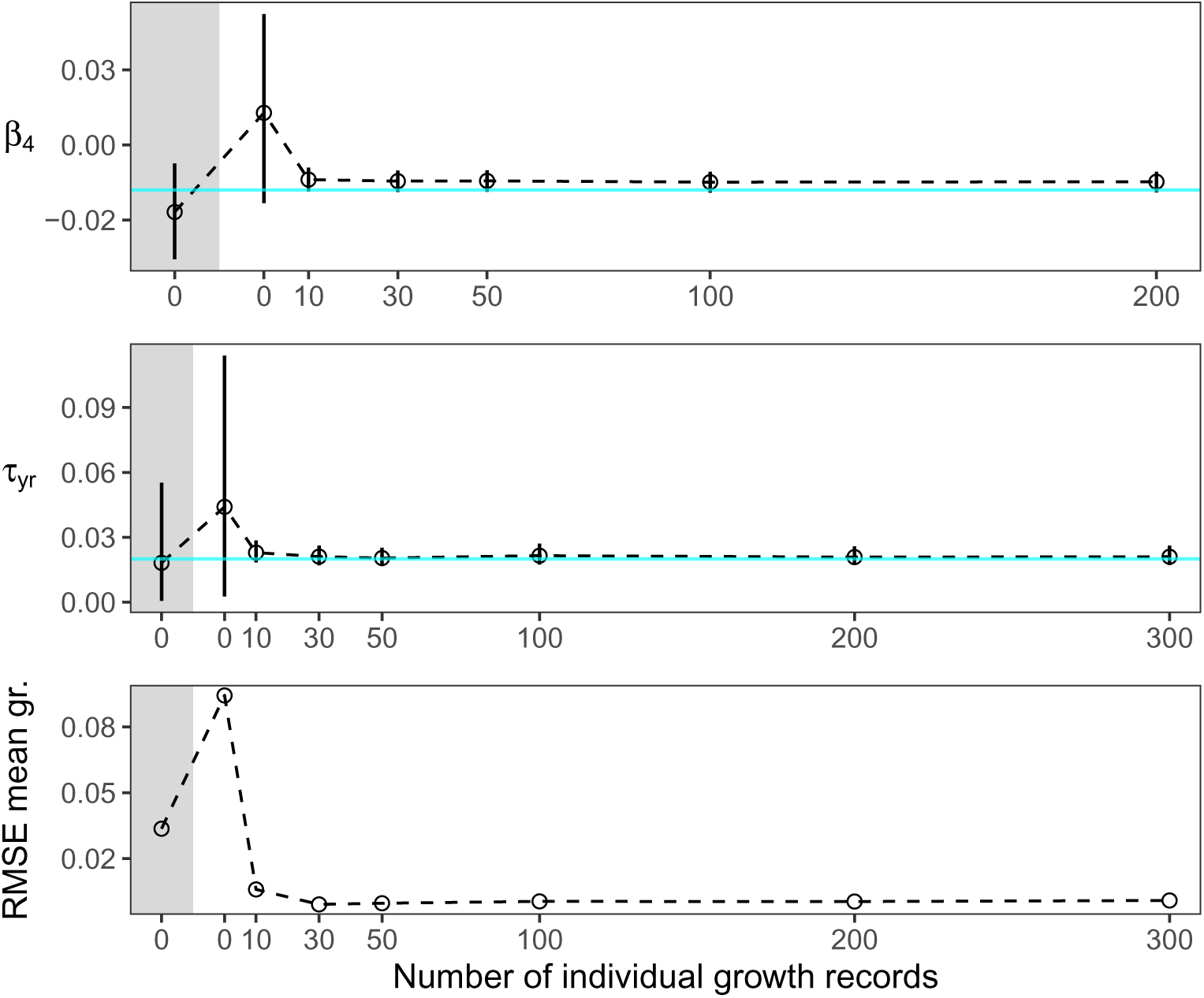
Posterior summaries of annual growth effect parameters including the coefficient for vapor pressure deficit *β*_4_ (a) and annual growth random effect standard deviation *τ*_yr_ (b) relative to simulated parameter values, along with root mean square error for mean size-structured growth rate (c) as a function of the number of individual growth records used to fit the model (population model was applied for 0 individual records). Grey shading indicates that annual rather than periodic population density data was applied (annual population model). Points and error bars in (a) and (b) represent posterior means and associated 95 percent credible intervals with cyan line indicating the simulated parameter value.

**Figure 4:**
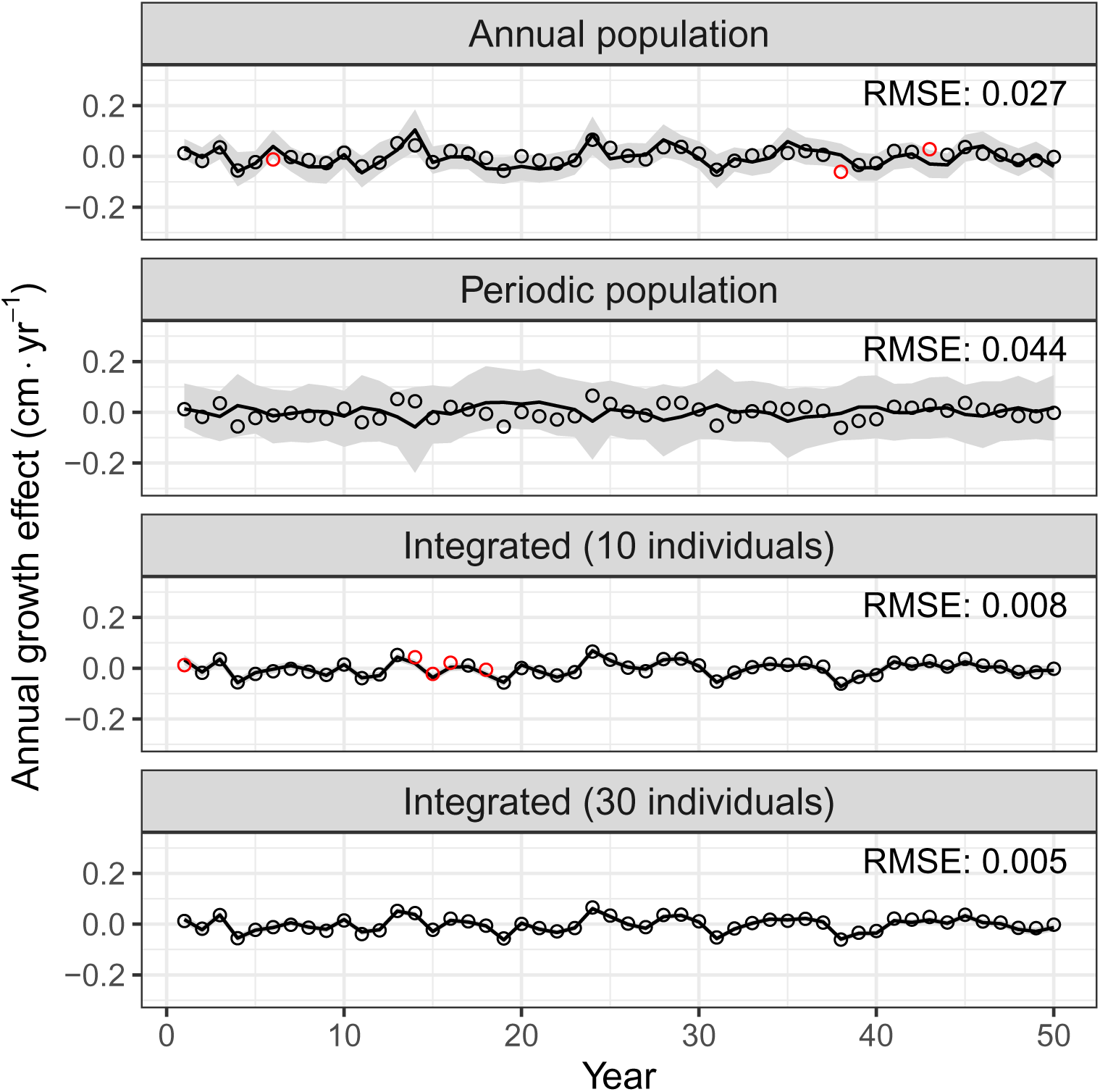
Posterior estimates of simulated annual growth effects (calculated as the sum of weather and annual random effects as defined in Table 1) under different simulated data scenarios. Lines and shading represent the posterior mean and associated 95 percent credible interval while points indicate simulated growth effect values. Points are colored red if they fall outside of the 95 percent credible interval and black if they fall within the interval bounds.

The integrated model resulted in improved inference for weather and annual random effects relative to both population model scenarios (annual, periodic) for any number of individual growth records. Posterior inference for the VPD growth coefficient (*β*_4_) was nearly identical across all integrated model scenarios (Fig. 3a). Moving from 10 to 30 records improved posterior inference for the annual random effect standard deviation (*τ*_yr_) contributing to a slight reduction in the RMSE of mean size-structured growth with no change in inference when more than 30 growth records were applied (Fig. 3b,c). Similar results were found comparing the ability of the integrated model to estimate annual growth effects. Estimates improved slightly (greater accuracy, less uncertainty, lower RMSE) when moving from 10 to 30 individual growth records, but remained largely unchanged when additional records were integrated within the model (Fig. 4). In particular, with 10 records there were 5 years in which the integrated model failed to capture the simulated annual growth effect (as determined by intersection with 95 percent credible intervals), while with 30 or more records annual growth effects were captured in all model years (Fig. 4).

### 3.2 Birch Lake red pine site

Predictions of size-structured population density were similar for the population and integrated models applied to Birch Lake with a negligible difference in the mean lppd score between the two models (Table 2). This near equivalence was further reflected in predictions of total density for which there were minor differences in the RMSE of trees (*<* 2 trees) and basal area (*<* 1 m^2^) per ha (Table 2). Consistent with these results, the integrated and population models had similar mean size-structured population density estimates across the model period with no systematic differences from each other or the mean observed size distribution across forest inventory years (Fig. 5). Similar mean density estimates for the two models yielded similar predictions of size structured population density, trees per ha, and basal area per ha over the model period, which closely matched inventory observations and showed no signs of systematic error (Figs. S4, S5).

**Figure 5:**
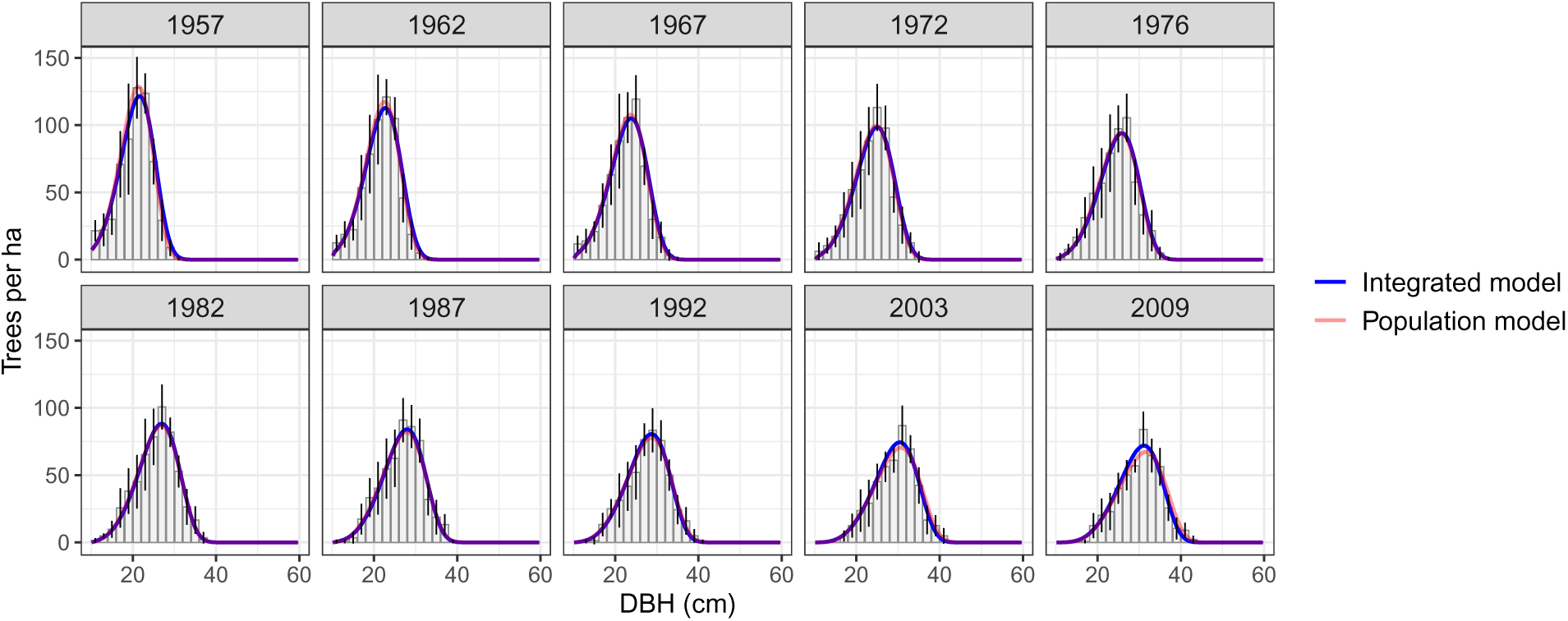
Mean size-structured population density estimates for models fit using population density and individual growth data (integrated model) and population density data alone (population model) in Birch Lake forest inventory years compared to mean inventory-plot-level counts within 2.0 cm diameter at breast height (DBH) size classes expressed on a per hectare scale. Error bars indicate two times the plot-level standard error in a given size class.

**Table 2:** Mean log pointwise predictive density (lppd) and root mean square error (RMSE) values for population density, individual growth rate, trees per hectare (TPH), and basal area per hectare (BAH) for models fit using both population density and individual growth data (integrated model) and population density data alone (population model) under 3-fold cross-validation for Birch Lake.

| Data | lppd |  | RMSE |  |
| --- | --- | --- | --- | --- |
|  | Population density | Individual growth | TPH | BAH |
| Integrated model | -2812.0 | 19955.8 | 156.4 | 4.8 |
| Population model | -2804.7 | 14184.4 | 154.6 | 4.6 |

Contrary to population density predictions, there were large differences in the ability of the population and integrated IPMs to predict individual growth rates. The integrated model had a much higher mean lppd score than the population model indicating better predictive performance for individual tree growth rates (Table 2). Mean size-structured growth rate estimates (*δ_t_*(*x*)) under the integrated model closely approximated mean observed growth rates within modeled size classes with no signs of bias (Fig. 6a). Mean growth rate estimates under the population model had greater variability around observed values with evidence of positive bias particularly in years with high VPD (Fig. 6b).

**Figure 6:**
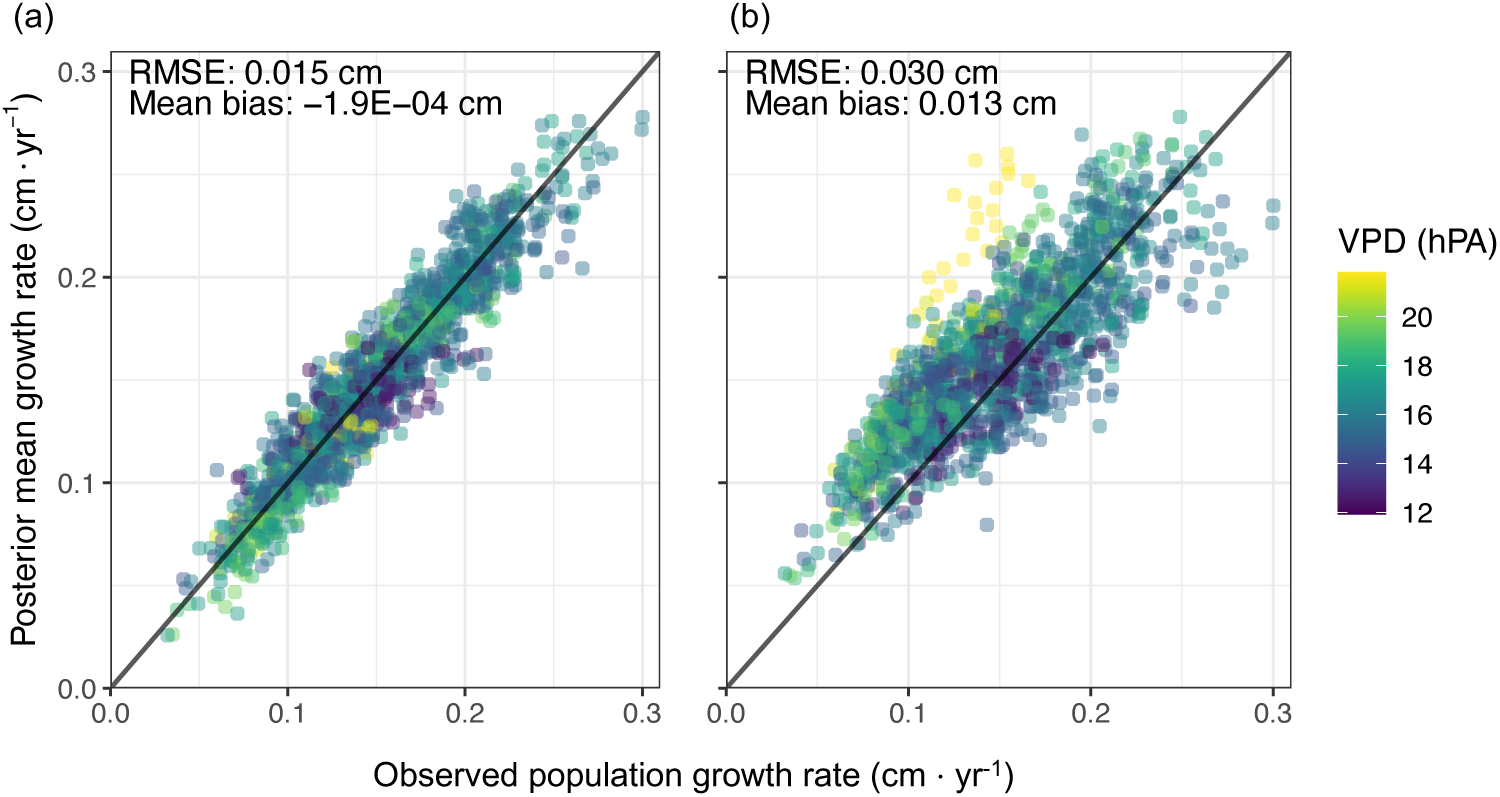
Mean size-structured growth rate (cm yr*^−^*^1^) estimates for Birch Lake applying the integrated model (a) and population model (b) plotted against mean observed growth rate within 0.3 cm size classes derived from individual growth records over the model period. Points are colored according to annual mean monthly vapor pressure deficit (VPD) in summer months (June-August).

Weather and annual random effect parameters were not identifiable under the population model. Specifically, MCMC chains had poor convergence (*R̂* > 1.05) for these parameters and approximately 30 percent of post burn-in iterations ended in a divergent transition. As a result, these parameters were dropped from the population model applied to the Birch Lake data (all presented results are for population model without weather and annual random effects). Shared growth parameters in the population and integrated models included size effect and population density effect parameters. Shared growth parameter estimates were similar between the two models with overlapping posterior distributions for all terms (Table S3, Fig. 7). On average, the relationship between size and mean growth was positive and concave, while the relationship between population density and mean growth was negative under both the population and integrated models (Table S3, Fig. S6). The magnitude and variation of size effect coefficients were lower under the integrated model (Figs. 7, S6).

**Figure 7:**
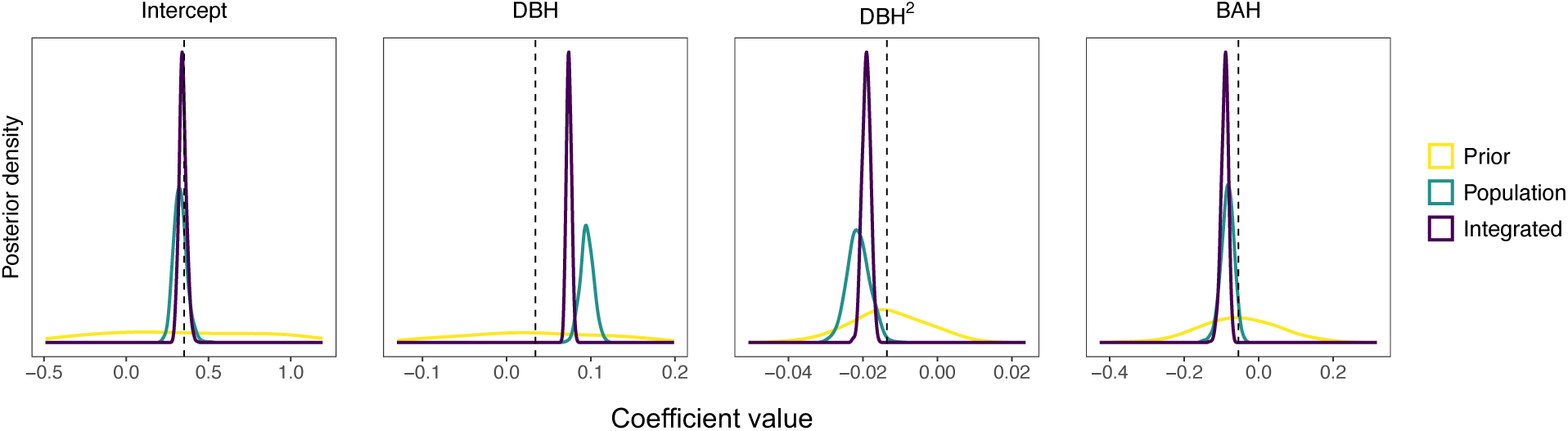
Posterior distribution of shared growth parameters for Birch Lake (Intercept, diameter at breast height (DBH), DBH^2^, and basal area per hectare (BAH)) under the integrated model applying both population density and individual growth data and the population model applying population density data alone relative to prior distributions. Dashed vertical lines indicate the prior mean parameter value.

Annual growth effects (equal to the sum of weather and annual random effects) included in the integrated model ranged between 0.08 cm yr*^−^*^1^ (Fig. S7) and accounted for 26 percent of mean size-structured growth variation (95 percent credible interval (CI): 24-30 percent). Weather variables including summer VPD and maximum spring temperature explained 29 percent (95 percent CI: 2-47 percent) of the annual growth effect variation (Fig. S8). Current year and lagged VPD had negative associations, while maximum spring temperature had a positive association with mean annual growth (Table S3). The magnitude of weather coefficients, however, was relatively small with VPD having the largest value: approximately 0.02 cm yr*^−^*^1^ reduction in growth per one unit increase of VPD measured in hPA (Table S3). The remaining variability in annual growth effects was driven by the annual random effect, which had an estimated standard deviation (*τ*_yr_) of 0.02 cm *·* yr*^−^*^1^ (Table S3).

## 4 Discussion

The presented dynamical IPM integrates population density and individual growth rate data to predict size-structured population dynamics while improving inference on the factors affecting mean growth rates. Across applications to both the simulated and Birch Lake datasets, the population and integrated dynamical IPMs generated functionally equivalent predictions of size-structured population density. However, estimates of the underlying mean size-structured growth rate differed under the two models with the integrated model generating more accurate estimates for both the simulated (Fig. 3c) and Birch Lake datasets (Table 2, Fig. 6). Effectively, the population data provided to both dynamical IPMs constrained estimates of the latent size-structured density, but without individual growth rate data from which to learn, the population model did not accurately identify the factors driving changes in size-structured density over time. These results illustrate the potential for inversely calibrated IPMs to correctly predict population density while incorrectly identifying underlying demographic rates noted in previous studies (Ellner, 2012; González et al., 2016; Plard et al., 2019b).

Differences in mean size-structured growth rate estimates were driven primarily by weather and annual random effects, which were poorly identified under the population model. In the simulation analysis, informed priors for these parameters reduced identifiability issues, but posterior parameter estimates had large uncertainty under both population model scenarios and were biased under the periodic population model (Figs. 3, 4). In the Birch Lake analysis, weather and annual random effects were not identifiable under the population model and were only included in the integrated model in which they accounted for approximately 26 percent of the variation in mean growth rates over time. Missing weather effects under the population model led to overly high estimates of mean growth during dry years defined by large VPD (Fig. 6). Larger size effect coefficients under the population model (Fig. 7) further contributed to the slight positive bias in mean growth rate estimates.

Without accurate and precise estimates of the factors driving demographic rates, predictions of future size-structured density (for which no data exist) may be biased and/or have too large uncertainty to be useful. The risk of poor predictions is exacerbated when the demographic factors missed by the IPM are expected to change under future conditions (Fig. 1). This is the case in the current application. The weather effects that the population IPM either gets wrong (simulated data) or misses completely (Birch Lake data) relate to summer VPD and maximum spring temperature. VPD had the largest effect among the included weather variables and its exclusion in the population model for Birch Lake contributed to positive bias in mean growth rate estimates during known drought years (1961, 1988) with high VPD (Fig. 6) previously found to impact forest growth dynamics at the site (D’Amato et al., 2013). The mean and variance of VPD are expected to increase globally under future climate scenarios fundamentally altering forest population dynamics (McDowell et al., 2020). Well-calibrated IPMs that include such temporally variable environmental factors should provide more meaningful predictions of structured population dynamics under global change (Yates et al., 2018) as highlighted in Figure 1.

Integration of individual growth rate data resolved the identifiability issues described above and improved estimates of mean size-structured growth. Specifically, estimates of mean growth rate were less variable and more closely matched observed individual growth rates in the Birch Lake analysis (Fig. 6, Table 2). Results of the simulation analysis indicated that integrating data from as few as 10 individuals was enough to accurately estimate mean growth rate parameters with small reductions in uncertainty when integrating data from 30 individuals (Figs. 3, 4). There were diminishing returns in terms of improved accuracy and reduced uncertainty of mean growth rate estimates when data from more than 30 individuals was integrated within the dynamical IPM. While the annual population scenario improved estimates of mean growth rate parameters relative to the periodic population scenario, these estimates were poorer than those resulting from any integrated model scenario (Fig. 3). This suggests that it is not the temporal resolution of population density data that limits inference on mean annual growth rate parameters. Rather, individual growth rate data provides unique information to support inference on the factors affecting growth. Population density data (whether periodic or annual) does not contain the same level of information because modeled demographic processes (growth, mortality, regeneration) jointly affect changes in population density over time (Ellner, 2012).

Plard et al. (2019b) found similar benefits of applying an integrated dynamical IPM as those described here, namely accurate estimates of the environmental factors driving demographic rates resulting in improved prediction of structured population density. In that study, the integrated dynamical IPM was applied to population density and individual demographic data observed at the same annual temporal resolution with individual variability accounted for by a single continuous trait. The presented integrated dynamical IPM accommodates misaligned population and individual data and accounts for individual differences attributable to observed (size) and unobserved variables (individual random effect). These are important steps toward broader application of integrated IPMs given both the prevalence of these data issues and their potential to affect inference when integrating population and individual data (Pacifici et al., 2019). Additional extensions to increase the utility of the integrated dynamical IPM to predict structured population dynamics under global change include scaling the model to multiple species and locations. The challenge is the high-dimensionality of the latent structured density when modeling multiple species and locations as highlighted in Gelfand et al. (2013). Previous statistical research on low dimensional approximation of complex spatio-temporal processes provides a foundation for such extensions (Wikle and Hooten, 2010), however future work is needed to adapt these methods to multiple species and traits and connect them to underlying demographic processes.

The integrated dynamical IPM presented here synthesizes population density and individual demographic data to better estimate mean demographic responses to variable environmental conditions facilitating improved prediction of size-structured population dynamics under global change. While population density data is widely available, individual demographic data is more limited given additional time and effort required to collect it. Fortunately, study results suggest that a small number of individual records are sufficient to improve inference on the demographic factors underlying changes in structured population density. These results combined with the presented dynamical IPM framework to integrate disparate population and individual datasets represent an important advancement in the application of IPMs to predict structured population dynamics in response to environmental change.

## Acknowledgements

We thank the USDA Forest Service Northern Research Station (FS-NRS) for establishing and maintaining the Birch Lake study, and Doug Kastendick in particular for leading data collection at the site. We further thank Anthony D’Amato for sharing Birch Lake tree ring data. This work was supported by funding from the USDA FS-NRS, 23-JV-11242305-080.

