## Supporting information for "Integration of individual and population data to improve predictions of structured population dynamics"

**Authors:** Malcolm S. Itter

### S1 Supplemental tables and figures

Table S1: Dynamical model notation key

| Notation | | Description |
|---|---|---|
| Indices | | |
| $p$ | | inventory plots ($p = 1, \ldots, P$) |
| $t$ | | years ($t = 1 \ldots, T$) |
| $j$ | | size classes ($j = 1, \ldots, k$) |
| $i$ | | cored trees ($i = 1, \ldots, n$) |
| $\ell$ | | interior integration points ($\ell = 1, \ldots, \mathcal{L}$) |
| Data | | |
| $y_{j,p,t}$ | | stem count by size class |
| $z_{i,t}$ | | annual diameter growth rate (cm) |
| $\lvert A \rvert$ | | inventory plot size (ha) |
| $\delta^*$ | | maximum diameter growth rate (cm) |
| $k^*$ | | maximum growth in terms of size classes |
| $L$, $U$ | | lower and upper size bounds (cm) |
| $h$ | | size class width (cm) |
| $\omega_j(\ell)$ | | Gauss-Legendre quadrature weights |
| Parameters | | |
| $\theta_{\text{ic}}$ | $\text{tph}_0$ | initial stem density per ha |
| | $\mu_0$ | shape of initial Weibull probability density function |
| | $\kappa_0$ | scale of initial Weibull probability density function |
| $\theta_{\text{gr}}$ | $\boldsymbol{\beta}$ | growth rate coefficients |
| | $u_t$ | annual growth random effect |
| | $\tau_{\text{yr}}$ | standard deviation of annual growth random effect |
| $\theta_{\text{mr}}$ | $\boldsymbol{\gamma}$ | survivorship rate coefficients |
| $\theta_{\text{ar1}}$ | $\rho$ | first order residual autoregression coefficient |
| | $\sigma$ | residual (white noise) growth rate standard deviation |
| $\theta_{\text{ind}}$ | $w_i$ | individual tree random effect |
| | $\tau_{\text{ind}}$ | standard deviation of individual random effect |
| $\theta_{\text{pop}}$ | $\upsilon_{p,t}$ | inventory plot random effect |
| | $\phi$ | temporal decay parameter of plot random effect |
| | $\tau_{\text{inv}}$ | standard deviation of plot random effect |
| | $\psi$ | negative binomial overdispersion parameter |
| Transformed parameters | | |
| $\lambda_{j,t}$ | | mean latent intensity |
| $\delta_t(x)$ | | mean size-structured growth rate |
| $\epsilon_{i,t}$ | | individual residual growth rate |
| $\xi_t(x)$ | | mean size-structured survivorship rate |
| $\mathbf{H}_t$ | | propagator matrix |
| $g_t(x \mid y)$ | | growth kernel |
| $G_{j,t}^{j'}$ | | integral of growth kernel |

Table S2: Root mean square error of total population density in terms of stems and basal area per ha for each simulated data scenario (mean relative error is provided in parentheses expressed as a percentage).

| Scenario | Stem density (stems $\cdot$ ha$^{-1}$) | Basal area (m$^2$ $\cdot$ ha$^{-1}$) |
|---|---|---|
| Population annual | 22.7 (1.5) | 0.64 (1.4) |
| Population periodic | 19.0 (2.1) | 1.15 (2.3) |
| Integrated (10 individuals) | 18.0 (1.9) | 1.11 (2.2) |
| Integrated (30 individuals) | 13.1 (1.4) | 0.83 (1.5) |
| Integrated (50 individuals) | 16.8 (1.9) | 1.05 (2.0) |
| Integrated (100 individuals) | 10.4 (1.1) | 0.59 (1.0) |
| Integrated (200 individuals) | 6.4 (0.7) | 0.32 (0.7) |

Table S3: Posterior summary of model parameters estimated conditional on both inventory and tree ring data (integrated model) and inventory data alone (population model). Posterior means are presented with 95 percent credible interval bounds in parentheses. NA values indicate that the parameter was not included under the corresponding model. Parameters are as defined in Table S1.

| Parameter | Integrated model | Population model |
|---|---|---|
| Growth | | |
| $\beta_0$ | 0.344 (0.305, 0.389) | 0.327 (0.26, 0.415) |
| $\beta_1$ | 0.0740 (0.0687, 0.0797) | 0.0954 (0.0804, 0.111) |
| $\beta_2$ | -0.0191 (-0.0212, -0.0169) | -0.0212 (-0.0269, -0.0155) |
| $\beta_3$ | -0.0912 (-0.113, -0.0745) | -0.0837 (-0.12, -0.0528) |
| $\beta_4$ | -9.48e-03 (-0.0161, -2.75e-03) | NA |
| $\beta_5$ | -6.19e-03 (-0.0126, 6.11e-04) | NA |
| $\beta_6$ | 3.79e-03 (-2.24e-03, 0.0104) | NA |
| $\tau_{\text{yr}}$ | 0.0225 (0.0182, 0.028) | NA |
| $\tau_{\text{ind}}$ | 0.0419 (0.0391, 0.0445) | NA |
| $\rho$ | 0.707 (0.698, 0.716) | NA |
| $\sigma$ | 0.0304 (0.0302, 0.0306) | 0.1 (fixed) |
| Survivorship | | |
| $\gamma_0$ | 6.71 (6.05, 7.41) | 6.73 (6.03, 7.48) |
| $\gamma_1$ | 1.64 (1.36, 1.95) | 1.78 (1.48, 2.12) |
| $\gamma_2$ | -0.645 (-0.826, -0.466) | -0.647 (-0.818, -0.458) |
| Observation error | | |
| $\tau_{\text{inv}}$ | 0.101 (0.0684, 0.147) | 0.101 (0.0646, 0.149) |
| $\phi$ | 82.8 (35.7, 1.53e+02) | 82.2 (35.1, 1.45e+02) |
| $\psi$ | 5.08 (4.15, 6.28) | 5.08 (4.16, 6.27) |
| Initial conditions | | |
| $\text{tph}_0$ | 1.24e+03 (1.16e+03, 1.33e+03) | 1.26e+03 (1.17e+03, 1.35e+03) |
| $\mu_0$ | 5.79 (5.64, 5.93) | 6.00 (5.84, 6.17) |
| $\kappa_0$ | 22.4 (22.2, 22.5) | 22.0 (21.7, 22.4) |

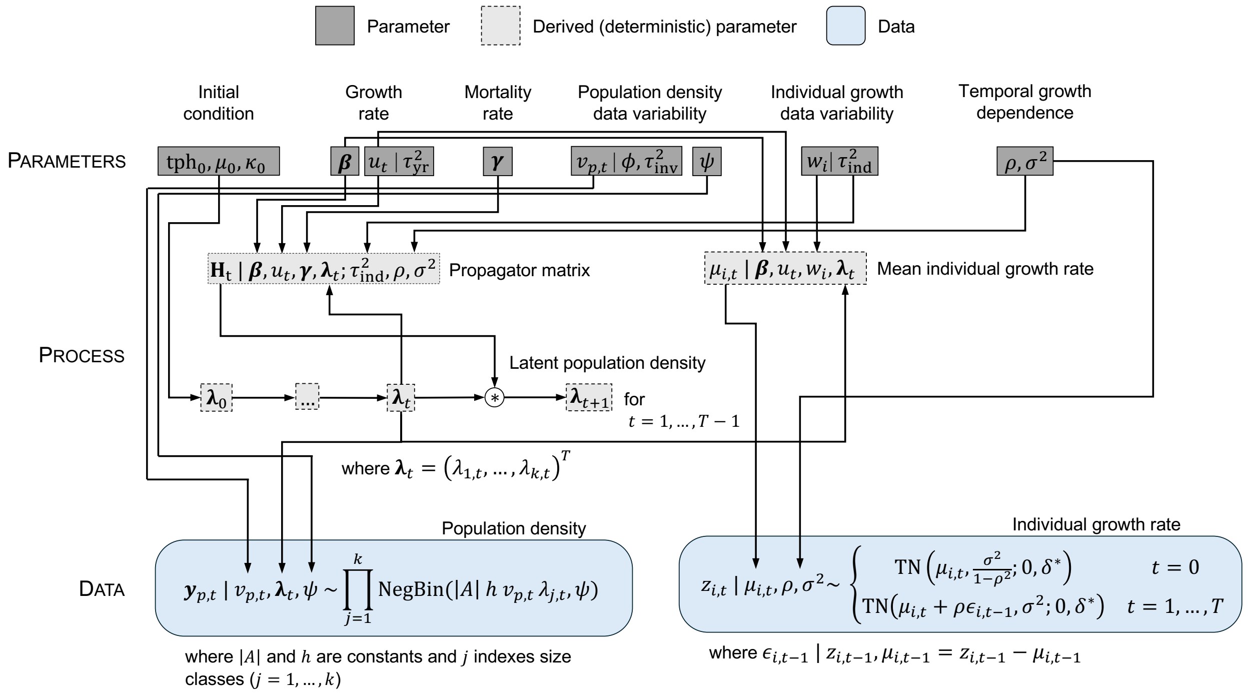


Figure S1: Directed acyclic graph for integrated dynamical integral projection model (IPM) synthesizing structured population density and individual growth rate observations.

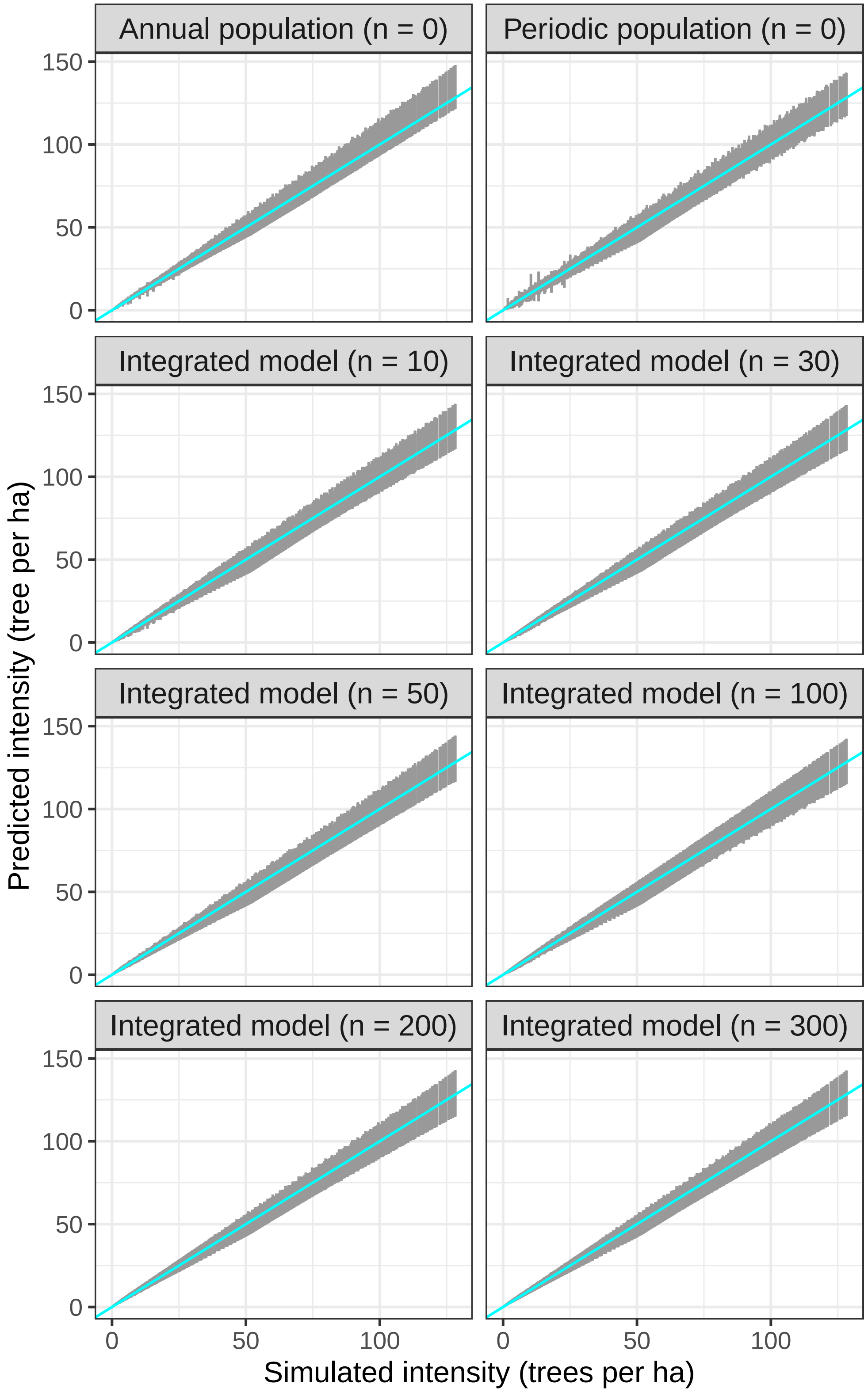


Figure S2: Posterior 95 percent credible intervals of latent stem density estimates versus simulated (true) values. Cyan line corresponds to a 1:1 relationship for reference.

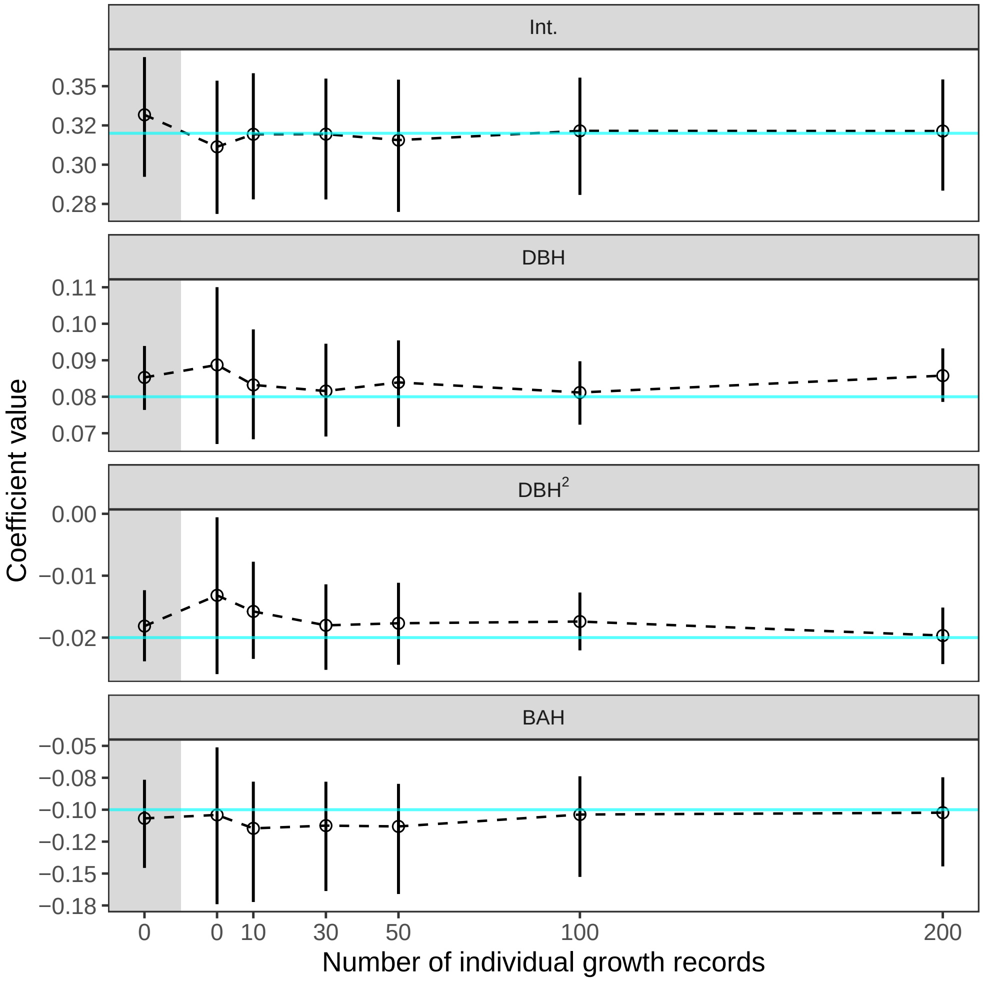


Figure S3: Posterior summaries for simulated mean size-structured growth parameters including the intercept and coefficients for diameter at breast height (DBH), $DBH^2$, and basal area per ha (BAH) as defined in Table 1 as a function of the number of individual growth records used to fit the model (population model was applied for 0 individual records). Grey shading indicates that annual rather than periodic population density was applied (annual population model). Points and error bars represent posterior means and associated 95 percent credible intervals with cyan line indicating the simulated parameter value.

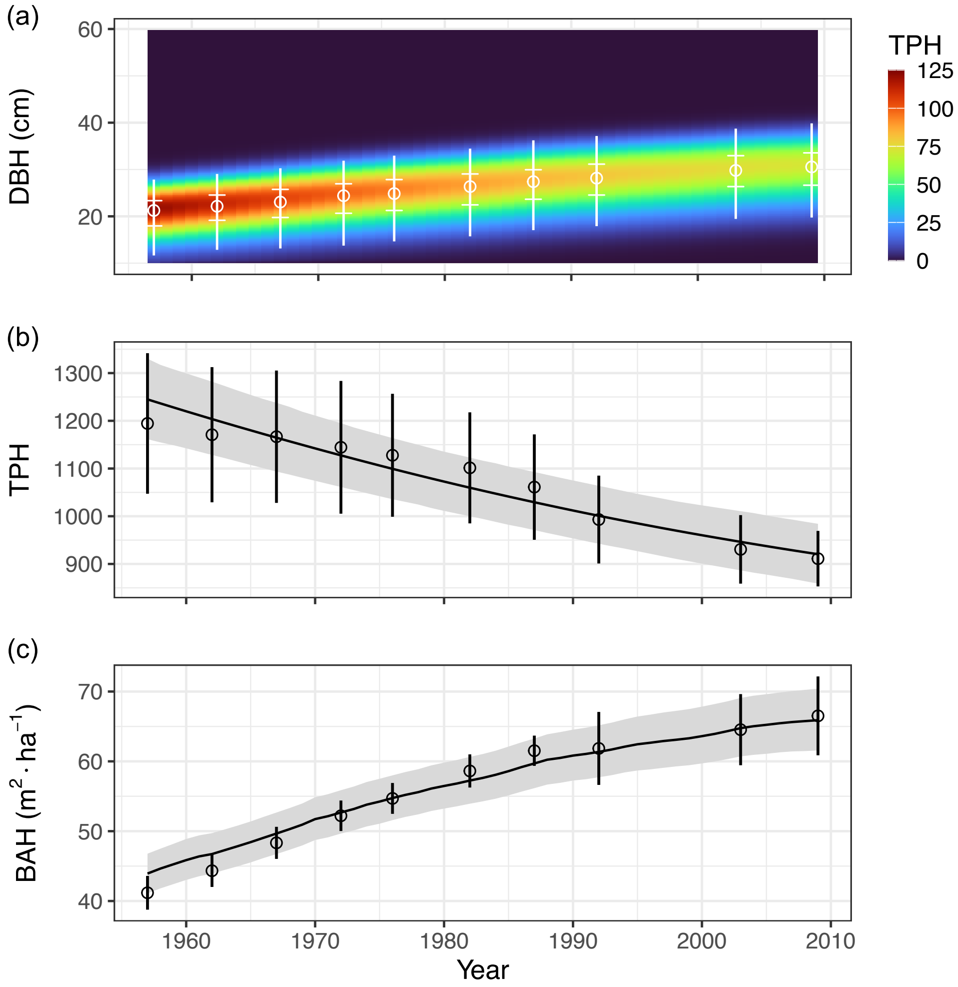


Figure S4: Predictions of size-structured population density (a), trees per ha (b), and basal area per ha (c) relative to forest inventory estimates applying integrated model to Birch Lake data. Points and tick marks in (a) correspond to the median and interquartile range of the mean observed size distribution with line bounds set to the 2.5 and 97.5 percentiles. Lines in (b) and (c) represent the posterior mean density estimate with corresponding 95 percent credible intervals shaded.

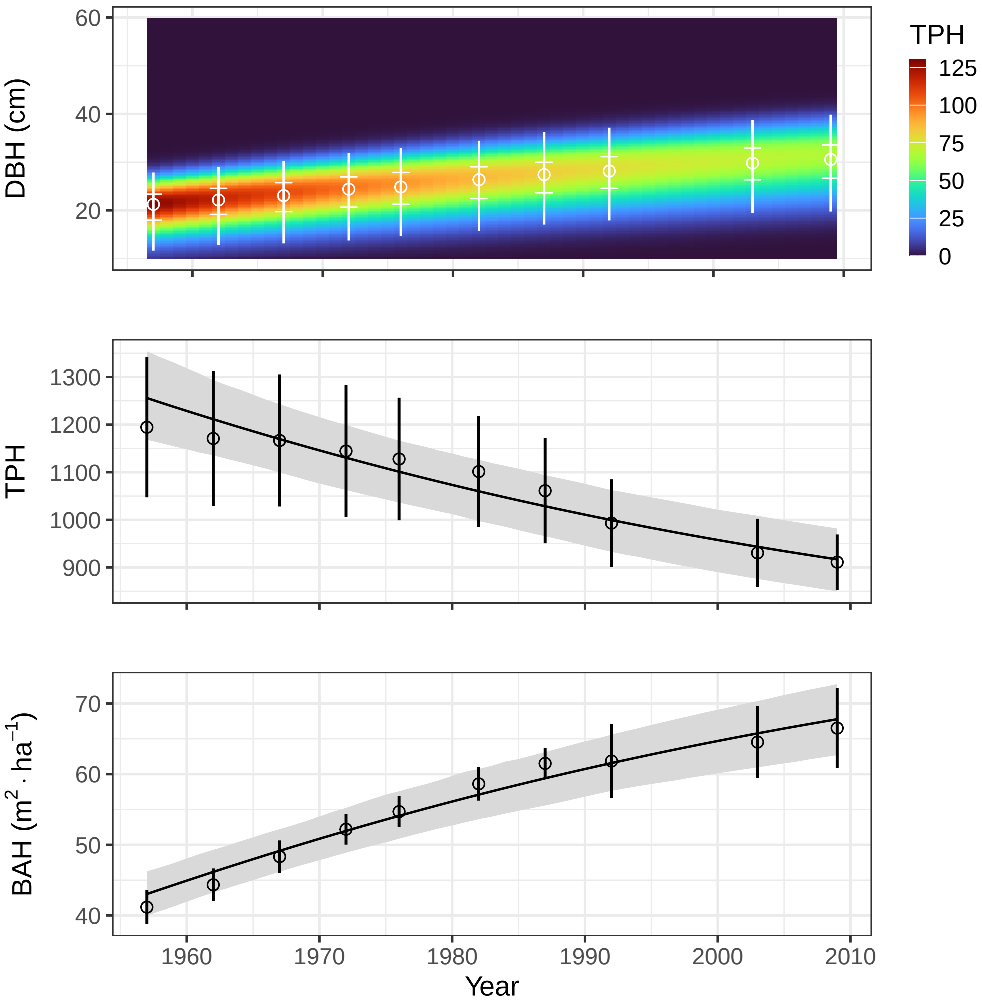


Figure S5: Predictions of size-structured population density (a), trees per ha (b), and basal area per ha (c) relative to forest inventory estimates applying population model to Birch Lake data. Points and tick marks in (a) correspond to the median and interquartile range of mean observed size distribution with line bounds set to the 2.5 and 97.5 percentiles. Lines in (b) and (c) represent the posterior mean density estimate with corresponding 95 percent credible interval shaded.

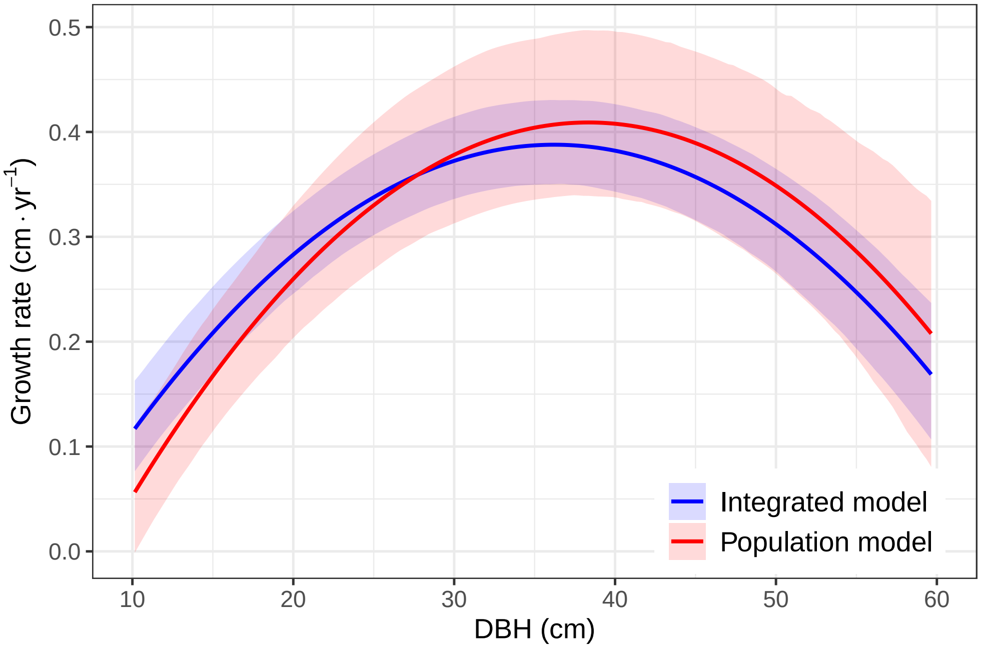


Figure S6: Predicted population growth rate versus tree size in terms of diameter at breast height (DBH). Predictions are made applying the intercept, DBH, DBH$^2$, and basal area per ha (BAH) terms under models fit using Birch Lake inventory and tree ring data (integrated) or inventory data alone (population). BAH was set to 35 m$^2$ per ha for all predictions. Lines indicate posterior means with shading representing 95 percent credible intervals.

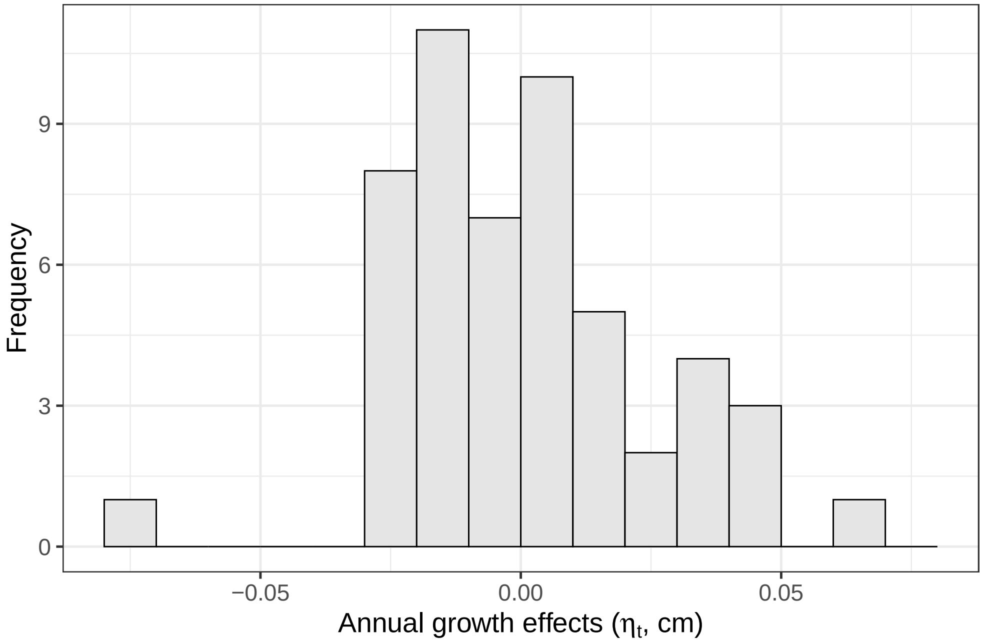


Figure S7: Histogram of posterior mean annual growth effects applying integrated dynamical model to the Birch Lake data.

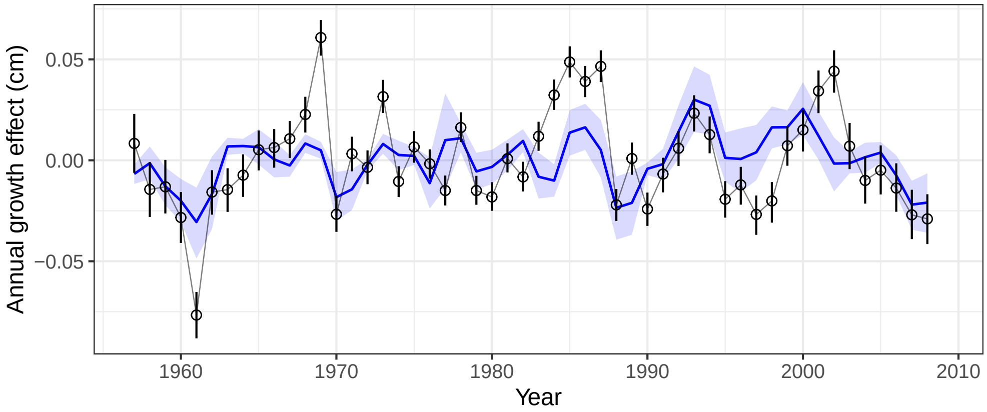


Figure S8: Posterior mean annual growth effects for Birch Lake site over time (points) along with posterior mean high-frequency growth effects predicted by weather variables (solid blue line). Uncertainty estimates correspond to 95 percent credible intervals represented as vertical lines (points) and shading (solid blue line).

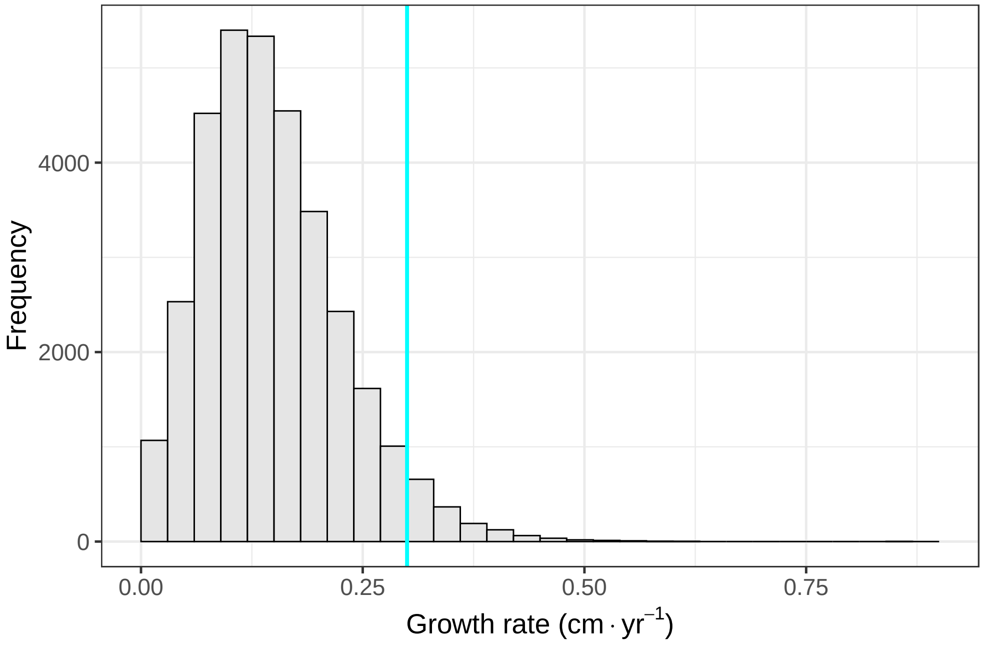


Figure S9: Histogram of observed annual diameter growth rates from Birch Lake tree ring data. Cyan line indicates the level of precision of inventory observations of tree diameter measured to the nearest 0.3 cm (0.1 in) with an English diameter tape.

### S2 Dynamical IPM implementation

#### S2.1 Numerical integration

The integral projection model (IPM) defined in Eqtn. (3) describes the temporal evolution of the latent intensity in continuous size ($s \in \mathcal{S} \subset \mathbb{R}^+$). The dynamical IPM involves three integrals over the continuous intensity in a given year $t$, one as part of the of the population data model (Eqtn. 1), another to approximate the basal area per hectare as a predictor of mean growth rate, and a third evaluation as part of the IPM process model (Eqtn. 3). All three integrals were solved numerically on a discrete scale applying the midpoint rule for the population data model and basal area per ha, and the bin-to-bin method for the IPM process model. The integral over the latent intensity is implicit in the presented negative binomial population data model (Eqtn. 1). Specifically, the observed tree sizes within an inventory plot $p$ in year $t$ represent a realization from an inhomogeneous Poisson point process with latent intensity $|A|\, v_{p,t}\, \lambda_t$. Discretizing the continuous intensity, the joint probability density for the observed size class counts is given by (adapted from Cressie, 1993),

$$\pi\left(y_{1,p,t}, \ldots, y_{k,p,t}, \sum_{j=1}^{k} y_{j,p,t}\right) \propto \exp\left(-\sum_{j=1}^{k} |A|\, h\, v_{p,t}\, \lambda_{j,t}\epsilon_{j,t}\right) \prod_{j=1}^{k} \left(\lambda_{j,t}\epsilon_{j,t}\right)^{y_{j,p,t}} \tag{S1}$$

where the summation term on the right-hand side represents the midpoint rule approximation of the integral of the continuous intensity and $\epsilon_{j,t}$ is a random effect term allowing for overdispersion of observed counts. Applying a Gamma prior for the random effect terms $\epsilon_{j,t} \sim \text{Gamma}(\psi^{-1}, \psi^{-1})$ and marginalizing over them leads to the negative binomial population data model presented in the main text (Lawless, 1987). We applied the same midpoint method to approximate the basal area per ha (bah) in the modeled forest at a given time $t$,

$$\text{bah}_t = \int_L^U \text{ba}(s)\lambda_t(s)\, ds \approx h \sum_{j}^{k} \text{ba}_j\, \lambda_{j,t}, \tag{S2}$$

where $\text{ba}_j$ is the basal area of the $j$th size class in $\text{m}^2$.

While we could apply the same midpoint method to approximate the integral in the IPM process model (Eqtn. 3), the mismatch in scale between inventory and tree ring observations necessitate a more robust approach. Specifically, inventory observations of tree size are measured to the nearest 0.3 cm (0.1 in) using diameter tape. The precision of the diameter tape sets the minimum size class that can be applied in the population data model. Applying a midpoint rule numerical integration approach with size classes set to the smallest observable size class ($h = 0.3$ cm) is problematic given that tree diameter growth rates are often much lower than this in closed canopy forests leading to an insufficiently fine integration mesh to accurately approximate modeled demographic rates (Zuidema et al., 2010). Indeed this was the case in the Birch Lake data where tree ring observations of annual diameter growth are small (generally $< 0.10$ cm per year) compared to coarse inventory observations of size (Fig.

S9). We applied a bin-to-bin numerical integration method to address this mismatch in scale (Dawson, 2013; Ellner et al., 2016).

Under the applied bin-to-bin method, the elements of the propagator matrix $\mathbf{H}_t$ describing the transition of the latent intensity from size class $j$ to $j'$ at time $t$ are given by,

$$\mathbf{H}_t[j', j] = \begin{cases} 0 & j' < j,\ j' > j + k^* \\ \frac{1}{h} \sum_{\ell}^{\mathcal{L}} \omega(\ell)\, \xi_{j,t}(\ell)\, G_{j,t}^{j'}(\ell) & j \leqslant j' \leqslant j + k^* \end{cases}, \tag{S3}$$

for $(j, j' = 1, \ldots, k)$. Here, $k^*$ is the maximum growth rate in terms of the number of size classes an individual can advance in a given year (Eqtn. S1), $\mathcal{L}$ is the number of integration points used to approximate the initial size distribution within size class $j$ indexed by $\ell$ $(\ell = 1, \ldots, \mathcal{L})$ with corresponding quadrature weights $\omega(\ell)$, $\xi_{j,t}(\ell)$ is the modeled survivorship of trees at integration point $\ell$ in size class $j$, and $G_{j,t}^{j'}(\ell)$ is the integral of the growth kernel expressing the transition from size class $j$ to $j'$ evaluated at point $\ell$. Under the applied truncated normal growth kernel, the inner integral (integrating over the initial size distribution) in the applied bin-to-bin method can be approximated applying the inverse normal cumulative density function (Dawson, 2013),

$$G_{j,t}^{j'}(\ell) = \frac{\Phi\left(x_{j'} + \frac{h}{2} | \mu_{j,t}(\ell), \nu^2\right) - \Phi\left(x_{j'} - \frac{h}{2} | \mu_{j,t}(\ell), \nu^2\right)}{\Phi\left(x_j - \frac{h}{2} + \delta^* | \mu_{j,t}(\ell), \nu^2\right) - \Phi\left(x_j - \frac{h}{2} | \mu_{j,t}(\ell), \nu^2\right)}, \tag{S4}$$

where $\Phi(x|\mu, \nu^2)$ indicates the cumulative normal density of $x$ given mean $\mu$ and variance $\nu^2$, $x_j$ is the midpoint of the $j$th size class, $\mu_{j,t}(\ell)$ is the mean growth rate evaluated at the $\ell$th initial size integration point for size class $j$, and $\nu^2$ is the variance of the truncated normal growth kernel. The upper truncation bound of the applied growth kernel is not strictly necessary (growth only needs to be constrained to be positive). We include it to minimize the number of normal cumulative density function evaluations $(2(k)(k + 1)$ without the upper truncation bound versus $4(k)(k^*)(\mathcal{L})$ with it) while restricting growth to a biologically reasonable range. The truncated normal growth kernel leads to a sparse banded propagator matrix $(\mathbf{H}_t)$ with non-zero elements along the diagonal and $k^* - 1$ sub-diagonals supporting efficient model implementation.

In both the Birch Lake and simulated data analyses we applied an $h = 0.3$ cm (0.1 in) step size to discretize the continuous intensity equivalent to the scale of DBH measurements. We applied a lower size bound of $L = 10$ cm (the smallest size recorded in forest inventory plots), and an upper size bound of $U = 60$ cm for the Birch Lake analysis and $U = 50$ cm for the simulated analysis selected to far exceed the maximum achievable size of individual trees during the model period to minimize the potential for eviction. The applied step size and lower and upper size bounds resulted in $k = 166$ size classes for the Birch Lake analysis and $k = 134$ size classes for the simulated analysis. In both analyses, we applied a maximum growth rate of $\delta^* = 1.8$ cm $\cdot$ yr$^{-1}$ set to exceed published maximum red pine growth rates for the Lake States (Bragg, 2001; Buckman et al., 2006). The maximum growth rate in terms of the number of size classes an individual tree can advance in a given year $(k^*)$ is calculated as the maximum growth rate divided by the discrete size class width rounded up to the nearest integer,

$$k^* = \left\lceil \frac{\delta^*}{h} \right\rceil, \tag{S5}$$

where $\lceil x \rceil$ denotes the ceiling function. We estimated a maximum growth rate equivalent to six size classes applying Eqtn. (S5) with $\delta^* = 1.8$ cm and $h = 0.3$ cm. Finally, we applied $\mathcal{L} = 4$ Gauss-Legendre integration points to approximate growth within each size class resulting step sizes of 0.02-0.10 cm, which closely approximate observed annual growth rates (inter-quartile range of observed growth rates: 0.09-0.19 cm).

#### S2.2 Demographic model selection

We applied inventory data from the USDA Forest Service Forest Inventory and Analysis (FIA) program to conduct variable selection for applied growth and survivorship functions in the Birch Lake analysis. Specifically, we used the rFIA package (Stanke et al., 2020; Doser et al., 2025) for the R statistical computing environment (R Core Team, 2024) to download and process FIA data for the Lake States including Minnesota, Wisconsin, and Michigan. We limited demographic analysis to FIA plots that had been measured at least two times, had a single condition class, and in which red pine represented at least 50 percent of the total basal area. We extracted individual tree information including diameter growth of live trees (trees that survived between two periodic inventories) and the size of mortality trees (trees that died between perioidic inventories) from the 'TREE_GRM_COMPONENT' table (USDA Forest Service, 2025). For the survivorship models, we removed all plots that had any disturbance event in order to accurately estimate natural mortality rates over time. Plots with disturbance were included in the growth models in order to ensure a wide range of growing conditions (e.g., forest density).

We conducted variable selection to identify the growth and survivorship models included within the dynamical IPM framework. We tested varying combinations of tree and forest density variables including diameter at breast height (DBH), $\text{DBH}^2$, and crown ratio at the tree scale, and basal area per ha, basal area per ha of trees larger than the target size class, relative stand density index, and realized to potential crown ratio based on the perfect plasticity approximation (Strigul et al., 2008) as measures of forest density. All variables were standardized by subtracting their mean and dividing by their standard deviation. We applied generalized linear models similar to standard IPM parameterization approaches (Rees et al., 2014). Growth was modeled applying an identity link while survivorship was modeled applying a logit link. We included a random effect for site in growth models to account for unobserved growth factors such as soil fertility. We initially included a random effect for site in survivorship models, but dropped it due to poor identifiability.

Candidate models were evaluated based on approximation of the leave-one-out cross-validation log pointwise predictive density estimated using the LOO R package (Vehtari et al., 2017). The selected growth model originally included tree crown ratio as an additional measure of size, however including this variable in the Birch Lake analysis led to a nearly singular design matrix due to limited variability in crown ratio values such that the variable was confounded with the intercept. As such, crown ratio was dropped from the version of the growth model included in the dynamical IPM. We also found that estimates of individual tree diameter growth had the best predictive performance for tree survivorship rates. However, we were not able to consistently identify the effect of modeled growth rate on survivorship within the dynamical IPM and so we applied the next best predicting survivorship model.

#### S2.3 Simulated data

Simulated data was generated from the proposed dynamical IPM framework setting parameters listed in Table S1 at fixed values. A single weather variable ($v$) was simulated to approximate vapor pressure deficit (VPD) over the 50-year simulated data period. We simulated $v$ as a weakly autoregressive process with normally distributed white noise errors as follows,

$$\begin{cases} v_0 \sim \mathrm{N}(0, \tau_v^2) \\ v_t \sim \mathrm{N}(\rho_v \, v_{t-1}, \tau_v^2) \qquad t = 1, \ldots, T \end{cases} \tag{S6}$$

with $\tau_v = 2$ and $\rho_v = 0.2$. The simulated high-frequency variable was provided as an input when fitting the dynamical IPM to simulated data. All other model components are as defined in the main text.

#### S2.4 Parameter models

Multiplicative plot random effects ($v_{p,t}$) were modeled independently across plots, but dependently over time applying a Gaussian process prior,

$$\mathbf{v}_p \sim \mathrm{N}\left(\mathbf{0}, \tau_{\mathrm{inv}}^2 C(\phi)\right) \tag{S7}$$

where $\mathbf{v}_p$ is the collection of random effects for plot $p$ over all inventory years, $\tau_{\mathrm{inv}}^2$ is the inter-annual variance in inventory observations, and $C(\phi)$ is an exponential temporal correlation matrix modeled as a function of the number of years between inventory measurements and temporal decay parameter $\phi$.

Individual growth random effects ($w_i$) are estimated applying a normal prior conditional on the individual growth variance term ($\tau_{\mathrm{ind}}^2$)

$$w_i \,|\, \tau_{\mathrm{ind}}^2 \sim \mathrm{N}(0, \tau_{\mathrm{ind}}^2). \tag{S8}$$

Annual growth rate random effect ($u_t$) are similarly estimated applying a normal prior conditiona on the inter-annual growth variance term ($\tau_{\mathrm{yr}}^2$),

$$u_t \,|\, \tau_{\mathrm{yr}}^2 \sim \mathrm{N}(0, \tau_{\mathrm{yr}}^2). \tag{S9}$$

We applied a standardized Weibull probability density function to approximate the initial condition

$$\lambda_0(j) = h^{-1}\,\mathrm{tph}_0 \left[ \frac{\pi\left(x_j|\mu_0,\kappa_0\right)}{\sum_j^k \pi\left(x_j|\mu_0,\kappa_0\right)} \right], \tag{S10}$$

where $\mathrm{tph}_0$ is the initial population density, $\pi$ is a Weibull probability density function conditional on scale $\mu_0$ and shape $\kappa_0$ parameters.

#### S2.5 Prior distributions

##### S2.5.1 Birch Lake analysis

*Demographic parameters*

Growth and survivorship parameters associated with size and population density were assigned informative priors to improve identifiability given that the dynamical IPM was fit to a single site. Parameter identifiability is a known issue with inverse calibration approaches for IPMs (González et al., 2016), and informed priors have been used in the past to address the data limitations that contribute to this issue (White et al., 2016). Growth $(\beta_0, \ldots, \beta_3)$ and survivorship $(\gamma_0, \gamma_1, \gamma_2)$ coefficients were assigned normal priors with hyperparameters set to the posterior mean and standard deviation from the prior demographic models fitted to independent FIA data.

$$\begin{aligned} \beta_j &\sim \mathrm{N}(\hat{\beta}_j, \hat{\sigma}^2_{\beta_j}) \\ \gamma_j &\sim \mathrm{N}(\hat{\gamma}_j, \hat{\sigma}^2_{\gamma_j}) \end{aligned} \tag{S11}$$

where $\hat{\beta}_j$, $\hat{\sigma}^2_{\beta_j}$ and $\hat{\gamma}_j$, $\hat{\sigma}^2_{\gamma_j}$ are the posterior mean and variance of the associated parameter from the prior growth and survivorship models fit to FIA data (section S2.2). Coefficients associated with weather variables used to estimate mean annual growth $(\beta_4, \beta_5, \beta_6)$ were assigned normal priors,

$$\beta_j \sim \mathrm{N}(0, 10^2) \tag{S12}$$

centered on zero with standard deviation set to match the scale of growth coefficients. Note that there was not a good way of setting informed priors for these coefficients since it is difficult to estimate annual growth effects using periodic FIA data alone. An important benefit of the dynamical IPM framework (or any inverse calibration approach for IPMs) is that it allows for estimation of correlation among demographic parameters through a joint prior distribution (González et al., 2016). We did not find evidence of strong correlation among modeled growth and survivorship parameters and thus applied independent priors for each demographic parameter.

*Growth variance and random effect parameters*

We assigned truncated normal priors to all growth standard deviation parameters with prior standard deviation of 0.5 to constrain parameters to the scale of annual growth rates.

$$
\begin{aligned}
\tau_{\text{ind}} &\sim \text{TN}_{[0,\infty]}(0, 0.5^2) \\
\tau_{\text{yr}} &\sim \text{TN}_{[0,\infty]}(0, 0.5^2) \\
\sigma &\sim \text{TN}_{[0,\infty]}(0, 0.5^2).
\end{aligned}
\tag{S13}
$$

Random effects ($\mathbf{u}$, $\mathbf{w}$) were sampled applying a non-centered parameterization,

$$
\begin{aligned}
u_t &= \tau_{\text{yr}}\, z_t \qquad t = (1, \ldots, T) \\
w_i &= \tau_{\text{ind}}\, z_i \qquad i = (1, \ldots, n).
\end{aligned}
\tag{S14}
$$

with $z_t$ and $z_i$ assigned standard normal priors. The AR1 coefficient for residual tree ring error was assigned a standard normal prior $\rho \sim \text{N}(0, 1)$ allowing for non-stationary errors.

*Inventory observation error parameters*

The standard deviation of the inventory plot random effect ($\tau_{\text{inv}}$) was assigned a truncated normal prior matching those for growth standard deviation parameters (Eqtn. S13). The associated temporal decay parameter ($\phi$) was also assigned a truncated normal prior with mean and variance set to reflect expected long-term temporal correlation (we expect a plot to remain above or below average conditions over the model period):

$$
\phi \sim \text{N}\left(\frac{3}{15}, 50^2\right). \tag{S15}
$$

Plot-level random effects were sampled applying a non-centered parameterization conditional on the Cholesky decomposition of the exponential covariance matrix ($\mathbf{L}$):

$$
v_{p,t} = \exp\left(\mathbf{L}\, z_{p,t}^{\text{inv}}\right), \tag{S16}
$$

with $z_{p,t}^{\text{inv}}$ assigned a standard normal prior ($p = 1, \ldots, P$, $t = 1, \ldots, T$). We applied a truncated normal prior to the square root of the inverse of the overdispersion parameter ($\psi$):

$$
\psi^{-\frac{1}{2}} \sim \text{TN}_{[0,\infty]}(0, 0.25^2). \tag{S17}
$$

*Initial condition parameters*

Initial condition parameters include the initial stem density ($\text{tph}_0$), and scale and shape parameters for the initial Weibull size distribution ($\mu_0$, $\kappa_0$). It was challenging to set meaningful priors for these parameters without initial information. Many models of forest dynamics apply a spin-up phase running the model to equilibrium to simulate initial conditions, but such methods do not apply to a dynamical model fit to on-the-ground conditions that may be far from equilibrium. To address this issue, we initialized the dynamical model in the first year that inventory data was collected using priors with hyperparameters estimated applying initial size and density observations. This initial inventory data was subsequently removed from the dynamical model fit (i.e., we used the initial inventory observations only to set hyperparameters for initial condition priors, not to fit the dynamical model, so the data was applied only once). This approach allows for uncertainty in inventory-based initial condition estimates while providing sufficient information to constrain initial conditions to a reasonable range.

All three initial condition parameters were assigned truncated normal priors with mean values set to match inventory-based estimates and standard deviations set to reflect prior uncertainty in these estimates,

$$\begin{aligned} \text{tph}_0 &\sim \text{TN}_{[0,\infty]}(\hat{\text{tph}}_0, 100^2) \\ \mu_0 &\sim \text{TN}_{[0,\infty]}(\hat{\mu}_0, 0.1^2) \\ \kappa_0 &\sim \text{TN}_{[0,\infty]}(\hat{\kappa}_0, 0.5^2), \end{aligned} \tag{S18}$$

where $\hat{\text{tph}}_0$, $\hat{\mu}_0$, $\hat{\kappa}_0$ are inventory-based estimates. We set $\hat{\text{tph}}_0$ to the mean trees per ha across inventory plots in the intitial inventory year and used method of moments estimators for Weibull scale and shape parameters (Justus et al., 1978),

$$\begin{aligned} \hat{\mu}_0 &= \left(\frac{\tau_0}{\hat{\bar{d}}_0}\right)^{-1.086} \\ \kappa_0 &= \frac{\hat{\bar{d}}_0}{\Gamma\left(1 + \frac{1}{\hat{\mu}_0}\right)}, \end{aligned} \tag{S19}$$

where $\hat{\bar{d}}_0$ is the initial quadratic mean diameter of individual trees and $\tau_0$ is an estimate of the spread of the initial diameter distribution.

##### S2.5.2 Simulated analysis

We applied the same prior distributions for the simulated data as we did for the Birch Lake data with adjustments to select hyperparameter values to center priors on the simulation parameter values. Specifically, we centered growth and survivorship coefficient values ($\boldsymbol{\beta}$, $\boldsymbol{\gamma}$) on simulated parameter values with small prior standard deviation (either 1.0 or 0.1 depending on the scale of the coefficient value). Note that the coefficient for the high frequency growth variable had an informed prior for the simulated data as opposed to uninformed priors for the Birch Lake data. Growth variance parameters and random effects were assigned the same priors as the Birch Lake analysis except the prior standard deviations for $\tau_{\text{yr}}$, $\tau_{\text{ind}}$, and $\sigma$ were reduced from 0.5 to 0.2 to more closely match simulated parameter values. Priors for inventory observation error parameters were unchanged. Priors for initial condition parameters were also kept the same except the prior means were set at simulated parameter values.

#### S2.6 Model implementation

For all applications of the dynamical IPM, three MCMC chains were run for a total of 2,000 iterations with a burn-in of 1,000 iterations. Model convergence was assessed based on R-hat values ($< 1.01$), bulk effective sample size (minimum of 100 per chain), and visual inspection of chains for all parameters (Vehtari et al., 2021).

#### S2.7 Starting values

Starting values for the Birch Lake and simulated analyses were sampled from prior distributions for all parameters under the inventory data only, population model fit. For the integrated model fit, we calculated residual growth rate values by subtracting estimated growth rates (applying starting values for growth rate parameters) from observed (or simulated) tree ring growth rates and then averaging by year or individual tree to estimate starting values for annual and individual random effects and associated parameters.

#### S2.8 Model evaluation

Mean log posterior predictive density (lppd) scores were approximated across Birch Lake cross-validation folds for both population density and individual tree growth data. All approximations of lppd were calculated following Gelman et al. (2014) and multiplied by -2 to be on the deviance scale. For population density, the mean lppd score was estimated as,

$$-2\left[\frac{1}{3}\sum_{m=1}^{3}\sum_{p=1}^{P_{\text{oos}}^{m}}\sum_{t=1}^{T_{\text{inv}}}\sum_{j=1}^{k}\log\left(\frac{1}{S}\sum_{s=1}^{S}\text{NegBin}\left(y_{j,p,t}\mid |A|\,h\,v_{p,t}^{(m,s)}\,\lambda_{j,t}^{(m,s)},\psi^{(m,s)}\right)\right)\right] \tag{S20}$$

where $m$ indexes the CV fold ($m = 1, \ldots, 3$), $s$ the posterior sample ($s = 1, \ldots, S$), and $P_{\text{oos}}^{m}$ is total number of held-out inventory plots under the $m$th CV fold. The $y_{p,t}(j)$ values are held-out size class observations and parameters to the right of | are sampled conditional on the training data for the $m$th fold. For individual tree growth, the mean lppd score was estimated as,

$$-2\left[\frac{1}{3}\sum_{m=1}^{3}\sum_{i=1}^{n_{\text{oos}}^{m}}\sum_{t=1}^{T_i}\log\left(\frac{1}{S}\sum_{s=1}^{S}\text{TN}_{[0,\,\delta^{*}]}\left(z_{i,t}\mid \delta_t(x_i)^{(m,s)}+w_i^{(m,s)}+\rho^{(m,s)}\,\epsilon_{i,t-1}^{(m,s)},\,\sigma^{2(m,s)}\right)\right)\right] \tag{S21}$$

where $n_{\text{oos}}^{m}$ is the number of held-out tree ring records under the $m$th CV fold, $z_{i,t}$ values are held-out individual annual growth rate observations, and the individual growth parameters are sampled using training data for the $m$th CV fold.

Root mean square error values are calculated as,

$$\left[\frac{1}{n}\sum_{i=1}^{n}(y_i - \hat{y}_i)^2\right]^{\frac{1}{2}} \tag{S22}$$

where $y_i$ is the observed value, $\hat{y}_i$ is the posterior mean estimated value, and $n$ is the total number of observations.

Variance partitioning was applied to estimate the proportion of annual growth rate variation attributable to annual growth effects and the proportion of annual growth effect variability explained by high-frequency weather variables. The proportion of annual growth rate variation attributable to annual growth effects was calculated following Schulz et al. (2025),

$$\frac{(Tk)^{-1}\sum_{t=1}^{T}\sum_{j=1}^{k}\left(\eta_t^{(s)} - \bar{\eta}^{(s)}\right)^2}{(Tk)^{-1}\sum_{t=1}^{T}\sum_{j=1}^{k}\left(\delta_{j,t}^{(s)} - \bar{\delta}^{(s)}\right)^2}, \tag{S23}$$

where $\eta_t$ is the annual growth effect equal to the sum of weather effects and the annual random effect ($\eta_t = \beta_4\,\mathrm{VPD}_t + \beta_5\,\mathrm{VPD}_{t-1} + \beta_6\,\mathrm{T}_t^{\mathrm{MAX}} + u_t$) and $\bar{\eta}$ is the mean annual growth effect over model years, $\bar{\delta}$ is the mean population growth rate over modeled size classes and years, and $s$ denotes the posterior sample ($s = 1, \ldots, S$). We calculated the Bayesian R-squared value (Gelman et al., 2019) to estimate the proportion of annual growth effect variability explained by high-frequency weather variables,

$$\frac{V_{t=1}^{T}\,\mathbf{x}_t'\boldsymbol{\beta}_w^{(s)}}{V_{t=1}^{T}\,\mathbf{x}_t'\boldsymbol{\beta}_w^{(s)} + \tau_{\mathrm{yr}}^{2\;(s)}}, \tag{S24}$$

where $\boldsymbol{\beta}_w$ is the collection of weather coefficients ($\beta_4, \beta_5, \beta_6$) and $V_{t=1}^{T}$ indicates the variance over model years.

##### S2.8.1 Model checking

In addition to out-of-sample model validation, we checked in-sample predictions for all Birch Lake model fits. We conducted visual posterior predictive model checks for population density based on inventory data and individual tree growth rates based on tree ring data. Further, we calculated Bayesian p-values for the mean, standard deviation, and chi-square statistics of size-structured population density across in-sample forest inventory plots for the three-fold cross-validation sets and the complete data application.
